# Time-of-flight–resolved interferometric speckle-contrast optical spectroscopy (TOF-iSCOS) for depth-resolved blood-flow sensing

**DOI:** 10.64898/2026.07.03.736163

**Authors:** Klaudia Nowacka-Pieszak, Marcin Marzejon, Neda Mogharari, Dawid Borycki

## Abstract

**Significance:** Continuous, noninvasive, and depth-resolved monitoring of blood-flow-related tissue dynamics remains an important unmet need. Speckle-contrast optical spectroscopy (SCOS), including interferometric implementations such as iSCOS, provides a scalable optical route to blood-flow sensing, but conventional continuous-wave approaches lack intrinsic depth selectivity. Time-of-flight (TOF) gating offers a way to separate superficial and deeper dynamic contributions in layered tissues, such as skin– muscle or scalp–cortex, by resolving photon path lengths.

**Aim:** We introduce a swept-source, single-channel implementation of interferometric speckle-contrast optical spectroscopy (iSCOS) to obtain TOF-resolved temporal speckle contrast, *κ*^2^, from the measured field autocorrelation *g*_1_, and evaluate its feasibility for depth-resolved blood-flow sensing.

**Approach:** A swept-source iNIRS system operating at 780 nm acquired interferometric signals, which were Fourier-transformed along the optical-frequency axis to recover complex TOF-resolved speckle fields. Temporal speckle contrast was then estimated at each TOF gate indirectly from *g*_1_ using the speckle-visibility relation. Diffusion-based numerical simulations were first used to compare the direct variance-based estimator and the indirect *g*_1_-based estimator under varying reduced scattering coefficient, diffusion coefficient, additive noise level, and bi-layer geometry. Because the simulations showed that the *g*_1_-derived *κ*^2^ estimator was substantially less sensitive to additive noise than the direct estimator, this estimator was used for the main phantom and *in vivo* analyses, while the direct estimator served as a simulation comparator. The *g*_1_-derived estimator was then applied to liquid and bi-layer phantoms, followed by proof-of-concept *in vivo* measurements on the human forearm during cuff occlusion and on the forehead during a Sudoku task.

**Results:** TOF-resolved *κ*^2^ curves recovered with the *g*_1_-derived estimator matched DWS theory across scattering coefficients, photon path lengths, and exposure times. The estimator preserved theoretical accuracy for additive noise amplitudes up to 50% of the field amplitude, whereas the direct variance estimator showed substantial noise-induced bias and required correction. Bi-layer simulations and phantom experiments reproduced the predicted direction and onset of TOF-dependent decorrelation-rate trends in layered media. In vivo, the recovered blood-flow index tracked the expected TOF-dependent cuff-occlusion and reactive-hyperemia response in the forearm. During the single-subject Sudoku task, the left-forehead recording showed a TOF-dependent relative blood-flow-index increase of +0.8 ± 1.9% at TOF = 400 ps, +9.8 ± 2.2% at TOF = 600 ps, and +15.2 ± 5.6% at TOF = 800 ps. This pattern is consistent with increased sensitivity to deeper tissue at longer photon path lengths, but requires cohort-level validation before quantitative interpretation as cognitive activation.

**Conclusions:** Coupling temporal speckle-contrast analysis with swept-source iNIRS yields a proof-of-concept, depth-resolved platform for blood-flow sensing. By estimating TOF-resolved speckle contrast through the *g*_1_-derived *κ*^2^ route, TOF-iSCOS suppresses additive-noise bias while preserving sensitivity to deeper dynamic tissue layers. The present single-channel results bridge continuous-wave iSCOS, interferometric NIRS and time-domain diffuse correlation spectroscopy (TD-DCS), and motivate future multi-channel and cohort studies for scalable cortical hemodynamic monitoring.

## 1. Introduction

Continuous and noninvasive assessment of cerebral blood flow (CBF) in the human brain remains an important goal in biomedical optics, neuroscience, and clinical physiology. CBF delivers oxygen and metabolic substrates to neural tissue, and its dynamic regulation is central to neurovascular coupling, autoregulation, functional activation, and brain health. Changes in CBF are therefore informative across a wide range of contexts, from cognitive and sensory stimulation studies to critical-care, perioperative, and neurological monitoring. In practice, however, resolving human cerebral blood-flow dynamics requires a demanding combination of temporal resolution, portability, noninvasiveness, and depth specificity that remains challenging for existing technologies.

Functional magnetic resonance imaging (fMRI) provides high-resolution mapping of hemodynamic changes coupled to cerebral blood flow [1], but it is not suited to bedside, wearable, or ambulatory monitoring. Optical methods provide a complementary route by exploiting the near-infrared transparency of biological tissue to monitor cerebral dynamics in real time. Near-infrared spectroscopy (NIRS) [2,3] tracks oxygenation-related changes, while diffuse correlation spectroscopy (DCS) [4,5] estimates blood-flow-related dynamics from the temporal autocorrelation of speckle intensity. These methods are portable, noninvasive, and well tolerated, making them attractive for functional and bedside monitoring. However, conventional continuous-wave (CW) implementations remain strongly affected by extracerebral tissues and provide limited intrinsic depth specificity.

Time-domain (TD) optical methods address depth specificity by resolving the photon time of flight (TOF). Time-domain NIRS [6], TD-DCS [7,8], and interferometric near-infrared spectroscopy (iNIRS) [9] resolve photon path lengths and can provide access to optical and dynamic properties. However, canonical TD-DCS and iNIRS architectures often rely on single-mode-fiber, single-element detection, which limits optical throughput and constrains acquisition speed; equally important, they offer limited control over the trade-off between signal-to-noise ratio (SNR), sampling rate, and effective TOF resolution, which becomes critical when functional dynamics span multiple time scales.

Highly parallel detection partially relieves this throughput limitation. Multi-element single-photon avalanche diode (SPAD) arrays deliver up to *N* = 1024 channels for parallel DCS [10,11], while parallel interferometric architectures using line-scan or two-dimensional detector arrays have reached *N* ≈ 512 detection channels in interferometric diffusing-wave spectroscopy [12] and *N* ≈ 10,000 channels in continuous-wave parallel interferometric NIRS, CW-πNIRS [13]. Parallelization improves SNR and enables fast hemodynamic monitoring, but it also produces large data streams and requires high-speed detection hardware that remains costly.

Speckle-contrast optical methods offer an alternative route to blood-flow sensing by relating the contrast measured over a finite integration time *T* to the underlying speckle decorrelation dynamics. Multi-exposure speckle-contrast approaches, including speckle visibility spectroscopy, speckle-contrast optical spectroscopy, and speckle-contrast optical tomography, have demonstrated robust blood-flow indices in deep-tissue geometries [14–23]. Recently, interferometric SCOS, iSCOS, embedded temporal speckle-contrast analysis in a CW parallel interferometric architecture, allowing both the field autocorrelation *g*_1_(*τ*_*d*_), conventionally used in DCS, and the speckle contrast *κ*^2^(*T*) to be obtained from the same interferometric dataset, with an explicit field-level pathway for correcting additive noise [24]. However, like all CW SCOS variants, this approach lacks intrinsic depth selectivity. Superficial tissues dominate the measured contrast and can mask deeper hemodynamics unless long source–collector separations are used. However, increasing the source–collector distance substantially attenuates the detected photon flux, thereby reducing signal-to-noise ratio, spatial resolution, and overall sensitivity.

Here we extend iSCOS into the time-of-flight-resolved regime, yielding TOF-iSCOS. To implement this concept experimentally, we used a swept-source iNIRS system in which a tunable laser sequentially samples the optical-frequency axis. This acquisition produces complex interferometric signals that encode path-length-dependent speckle dynamics. After Fourier transformation along the optical-frequency axis, these signals provide TOF-resolved complex representations of the speckle field. TOF-resolved temporal speckle contrast, *κ*^2^(*τ*_*s*_, *T*), can then be estimated in two ways: directly, from the variance-to-mean-squared ratio of the TOF-gated field fluctuations over a temporal integration window *T*; or indirectly, from the integral relation between the TOF-gated field autocorrelation, *g*_1_(*τ*_*s*_, *τ*_*d*_), and speckle visibility [16,20,25]. Here, *τ*_*s*_ denotes photon time of flight and *τ*_*d*_ denotes the correlation lag time.

In the present single-channel implementation, we used the indirect *g*_1_-based estimator of *κ*^2^(*τ*_*s*_, *T*) for the main phantom and *in vivo* analyses. This choice was motivated by numerical simulations showing that the *g*_1_-based estimator is substantially less sensitive to additive-noise bias than the direct variance-based estimator at the TOF-gate level. The direct estimator was therefore retained as a simulation comparator rather than used for the primary experimental analysis. Crucially, the same swept-source iNIRS data stream provides both the distribution of times of flight and the TOF-resolved dynamic information, without requiring an external timing module.

Although the TOF-iSCOS framework is formulated for parallel detection, current camera refresh rates remain one to two orders of magnitude below those required for finely resolved swept-source interferometric sampling, as comprehensively analysed in [26]. Therefore, in this proof-of-concept study we used a single-channel swept-source iNIRS system and relied on the autocorrelation-based relation between *g*_1_(*τ*_*s*_, *τ*_*d*_) and *κ*^2^(*τ*_*s*_, *T*) to obtain TOF-resolved temporal speckle contrast.

The overall TOF-iSCOS concept and workflow are summarized in Fig. 1. A swept-source iNIRS measurement first provides complex interferometric signals as a function of optical frequency. Fourier transformation along the optical-frequency axis then recovers TOF-resolved complex speckle fields and the corresponding distribution of times of flight. Within selected TOF gates, the field autocorrelation *g*_1_(*τ*_*s*_, *τ*_*d*_) is computed and converted into TOF-resolved temporal speckle contrast, *κ*^2^(*τ*_*s*_, *T*), using the speckle-visibility relation. This enables functional blood-flow readouts from early, intermediate, and late TOF gates, which preferentially weight superficial, mixed-depth, and deeper tissue dynamics, respectively.

**Figure 1.**
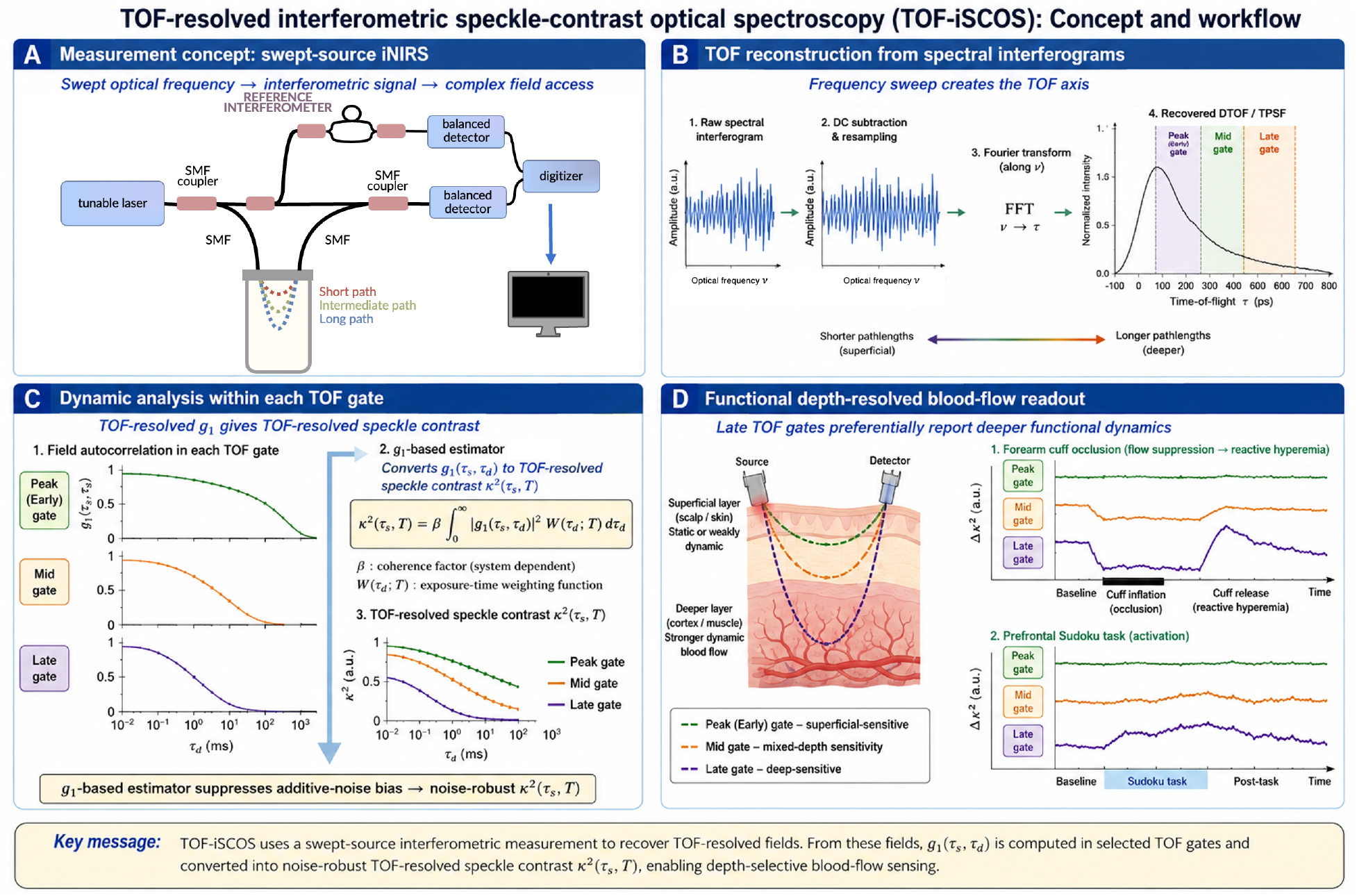
Concept and workflow of TOF-resolved interferometric speckle-contrast optical spectroscopy (TOF-iSCOS). (A) Single-channel swept-source iNIRS measurement concept. A tunable laser is swept in optical frequency, and the detected interferometric signal encodes the complex optical field from photons traveling along different path lengths. (B) After DC subtraction, resampling using the reference-interferometer signal, and Fourier transformation along the sweep axis, the time-of-flight distribution is recovered. Early, middle, and late TOF gates correspond to progressively longer photon path lengths and different depth sensitivities. (C) The TOF-gated field autocorrelation *g*_1_(*τ*_*s*_, *τ*_*d*_) is computed and converted into TOF-resolved temporal speckle contrast, *κ*^2^(*τ*_*s*_, *T*), using the speckle-visibility relation. This *g*_1_-based estimator improves robustness to additive noise compared with direct contrast estimation. (D) Early TOF gates preferentially sample superficial dynamics, whereas late TOF gates contain a higher contribution from deeper dynamic tissue, enabling depth-selective monitoring during vascular and functional activation paradigms.

We make four contributions. First, we derive the swept-source TOF-iSCOS forward model linking the TOF-resolved temporal speckle contrast, *κ*^2^(*τ*_*s*_, *T*), to the DWS-based TOF-dependent decorrelation rate, *ξ*(*τ*_*s*_). Second, we use numerical simulations spanning reduced scattering coefficient *µ*_*s*_′, diffusion coefficient, additive noise amplitude, and two-layer geometries to compare direct variance-based and indirect *g*_1_-based estimators of TOF-resolved temporal speckle contrast. These simulations show that the indirect estimator substantially suppresses additive-noise bias at the TOF-gate level and is therefore preferable for experimental analysis. Third, we apply this estimator to liquid and bi-layer phantoms using a swept-source iNIRS system operating at 780 nm, demonstrating TOF-resolved sensitivity to deeper scattering layers. Fourth, we provide functional in vivo demonstrations of TOF-iSCOS on the human forearm during cuff occlusion and on the left forehead during a Sudoku task, showing TOF-dependent relative blood-flow-index changes consistent with TOF-gated probing of deeper tissue dynamics.

## 2. Theory

We summarize the elements of the TOF-iSCOS forward model needed to interpret the experiments. The signal-formation, distribution-of-times-of-flight (DTOF), and DWS-based field-autocorrelation building blocks follow the iNIRS framework introduced in [9]. We use DTOF to denote the photon time-of-flight distribution in the theoretical description; the experimentally measured counterpart is also referred to as the temporal point-spread function (TPSF) in figures and captions. Unless stated otherwise, DTOF and TPSF denote the same TOF-resolved intensity distribution. We adopt the same notation and conventions established there (analytic complex TOF-resolved field, Wiener–Khintchine relation between the cross-spectral density and the mutual coherence function, and the in-medium wavenumber *k* = 2*πn*/*λ*_0_ used in the original iNIRS paper [9]). The derivation parallels that of CW-iSCOS [24], with the addition of an optical-frequency sweep and the consequent appearance of a TOF axis. The model is formulated for a general multi-pixel detector, in which *r* denotes the detector position and ⟨ · ⟩_*r*_ denotes averaging over detector pixels. In the experiments reported here, however, we used a single detector channel. Therefore, no spatial detector averaging was performed, and the expressions involving ⟨ · ⟩_*r*_ should be interpreted as their single-pixel limit. Throughout, *v* denotes optical frequency, *τ*_*s*_ the photon time of flight introduced by the Fourier transform with respect to *v, τ*_*d*_ the lag time of the field autocorrelation, and *T* the temporal integration window used for temporal speckle-contrast analysis. Quantities expressed in the optical-frequency (Fourier) domain are denoted with a tilde, whereas their corresponding TOF-domain representations are written without a tilde.

### 2.1 Swept-source TOF-iSCOS signal formation

In a swept-source Mach–Zehnder interferometer, the laser is tuned over a range Δ*v* of optical frequencies during the acquisition. At each frequency, the detector records the spectral interferograms

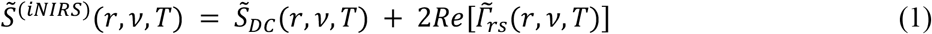

where the mutual coherence function

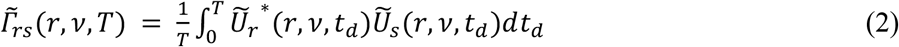

encodes the interference between reference and sample fields at exposure *T* in the Fourier- / frequency-domain. The DC offset is typically removed by mean subtraction, leaving the interferometric component

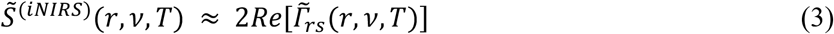

An inverse Fourier transform along *v* yields the time-of-flight–resolved quantity approximating the optical field scattered from the sample

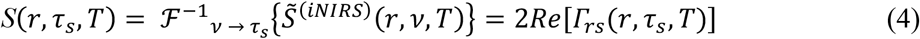

where *τ*_*s*_ is the photon time of flight associated with the optical path length distribution within the scattering medium and introduced by the wavelength-to-time pairing. Because 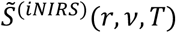 is real-valued, its inverse Fourier transform is Hermitian symmetric in *τ*_*s*_. Following the iNIRS framework [9,27], we discard the negative-*τ*_*s*_ mirror branch and interpret *S*(*r, τ*_*s*_, *T*) as the complex analytic TOF-resolved field on the positive-*τ*_*s*_ side.

The TOF-resolved interferometric field *S*(*r, τ*_*s*_, *T*) constitutes the fundamental observable of the TOF-iSCOS framework. Depending on the temporal acquisition strategy, it may be analyzed in two complementary ways.

When the exposure time *T* is sufficiently short relative to the underlying decorrelation dynamics, repeated acquisitions of *S*(*r, τ*_*s*_, *T*) enable direct estimation of the TOF-resolved temporal field autocorrelation function (here and below, *S*(*r, τ*_*s*_, *T*) denotes the complex analytic TOF-resolved field defined in Eq. (4), not the raw real-valued interferogram, so that the autocorrelation below is the first-order field autocorrelation rather than the autocorrelation of a real cosine)

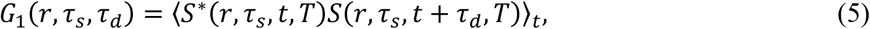

where ⟨ · ⟩_*t*_ denotes temporal averaging. The normalized autocorrelation is:

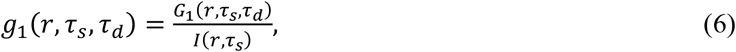

where *I*(*r, τ*_*s*_) = *G*_1_(*r, τ*_*s*_, *τ*_*d*_ → 0) denotes the TOF-resolved intensity distribution (distribution of times-of-flight, DTOF). Together, these quantities provide simultaneous access to both photon path-length distributions (*I*(*r, τ*_*s*_)) and depth-dependent flow dynamics (*g*_1_(*r, τ*_*s*_, *τ*_*d*_)) from the same interferometric dataset.

Integration over the TOF axis yields the conventional diffuse temporal autocorrelation function

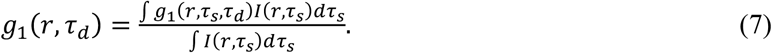

Alternatively, the exposure duration *T* itself may be varied and the temporal blurring of the fluctuating interferometric field analyzed through the TOF-resolved speckle visibility (contrast). Importantly, once sufficiently short-exposure measurements are available, longer effective exposure times may also be synthesized digitally through temporal averaging of consecutive interferometric measurements. Specifically, an effective exposure *T*_*m*_ = *N*_*m*_ *T*_0_ may be constructed from *N*_*m*_ consecutive short-exposure acquisitions with native exposure *T*_0_. This strategy is particularly advantageous when detector dynamic range is limited, as it reduces the risk of detector saturation and nonlinear response associated with long physical exposures. Furthermore, it preserves access to the underlying short-exposure data, allowing multiple effective exposure times to be generated retrospectively from a single acquisition.

The squared speckle contrast at a given TOF gate is defined as the variance of the TOF-resolved interferometric signal, normalized by the mean reference and sample intensities at that TOF:

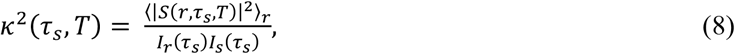

where *I* (*τ* ) = ⟨|*Ur,s* (*r,τ*_*s*_ ) |^2^⟩. Equation (8) is written for a general multi-pixel detector, where ⟨ · ⟩_*r*_ denotes averaging over independent detector realizations. In the present single-channel implementation, no spatial detector ensemble was available. Therefore, the experimental analyses used the autocorrelation-based estimator introduced below, while the direct variance-based estimator was evaluated only in numerical simulations.

Following the speckle-visibility-spectroscopy expansion [16], *κ*^2^(*τ*_*s*_, *T*) is related to the field autocorrelation by

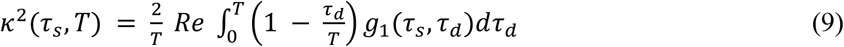

Equation (9) establishes the central connection underlying TOF-iSCOS. The TOF-resolved speckle contrast can be computed directly from the TOF-resolved temporal field autocorrelation function. Consequently, the same rapidly acquired interferometric dataset may be analyzed either through explicit *g*_1_(*τ*_*s*_, *τ*_*d*_) estimation or through exposure-dependent speckle visibility analysis without requiring separate measurement modalities. This relation unifies TOF-resolved iNIRS/DWS-style autocorrelation analysis and TOF-resolved SCOS/SVS contrast analysis within a common interferometric framework. In future multi-pixel implementations, the required ensemble averaging need not rely solely on temporal sampling. Instead, independent speckle realizations recorded simultaneously across detector pixels may be spatially combined, analogous to conventional SCOS, enabling robust TOF-resolved contrast estimation from a single camera exposure or a small number of exposures.

### 2.2 TOF-resolved temporal autocorrelation model

Within the diffusing-wave-spectroscopy framework, the TOF-resolved field autocorrelation is single-exponential,

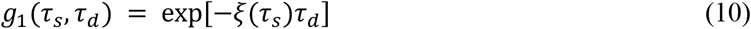

with the TOF-dependent decorrelation rate

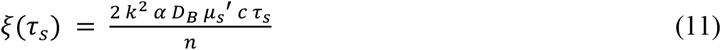

where *k* = 2*πn*/*λ*_0_ is the wavenumber of light in the sample (n the medium refractive index, *λ*_0_ the vacuum wavelength), following the iNIRS and DWS convention of [9,27], *αD*_*B*_ is the blood flow index, *µ*_*s*_′ the reduced scattering coefficient, *c* the speed of light, and *n* the medium refractive index. Substituting Eq. (10) into Eq. (9) and evaluating the integral analytically yields the closed-form expression

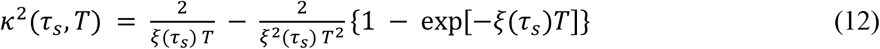

Equation (12) constitutes the central forward model of TOF-iSCOS speckle-visibility analysis. Three intuitive limits follow. For *T* ≪ 1/*ξ*(*τ*_*s*_), *κ*^2^(*τ*_*s*_, *T*) → 1 (frozen speckle). For *T* ≫ 1/*ξ*(*τ*_*s*_), *κ*^2^(*τ*_*s*_, *T*) → 2/[*ξ*(*τ*_*s*_)*T*] (speckle integrated to noise floor). The crossover scales linearly with *τ*_*s*_ through Eq. (12). Late-arriving photons sample longer optical paths and accumulate more dynamic phase, hence faster contrast decay.

### 2.3 Connection to the DTOF and depth selectivity

In a homogeneous semi-infinite medium, the distribution of times of flight *I*(*τ*_*s*_) follows the diffusion equation [28]. The non-TOF-resolved speckle contrast obtained without spectral sweeping is then a DTOF-weighted integral of Eq. (9),

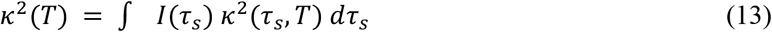

In Eq. (13) we adopt the convention that *κ*^2^(*τ*_*s*_, *T*) is the per-TOF squared contrast of Eq. (8), already normalized at each gate by the per-TOF mean intensities *I*_*r*_(*τ*_*s*_)*I*_*s*_(*τ*_*s*_), the DTOF *I*(*τ*_*s*_) then enters as a simple weight inside the integral, and the corresponding normalization ∫ *I*(*τ*_*s*_)*dτ*_*s*_ is absorbed in the per-TOF definition. When the non-TOF-resolved squared contrast is instead defined directly from the TOF-integrated interferometric signal, i.e. variance over [TOF-integrated mean reference intensity] × [TOF-integrated mean sample intensity], statistical independence of contributions from different TOFs (incoherent superposition of speckle fluctuations) yields the equivalent expression *κ*^2^(*T*) = [∫ *I*(*τ*_*s*_)^2^*κ*^2^(*τ*_*s*_, *T*)*dτ*_*s*_] / [∫ *I*(*τ*_*s*_) *dτ*_*s*_]^2^, with the squared DTOF weight in the numerator and the (∫ *Idτ*_*s*_)^2^ normalization in the denominator. The two conventions agree under our usage, and either may be used to compare TOF-iSCOS to non-gated CW iSCOS, clarifying why CW SCOS averages superficial and deep dynamics together. Restricting the integration in Eq. (13) to a TOF window [*τ*_*s*,1_, *τ*_*s*,2_] suppresses photons outside the gate and preserves the dynamics encoded in that subset of paths. In a layered medium, the late TOF window is enriched in photons that visited the deep layer. Consequently, *ξ*(*τ*_*s*_), measured in the late gate reports preferentially on deep dynamic scattering. We exploit this property in the bi-layer phantom and the *in vivo* experiments below.

### 2.4 Direct vs g_1_-based estimators

Two estimators of *κ*^2^(*τ*_*s*_, *T*) are useful in the TOF-iSCOS framework. The first is a direct estimator, in which the TOF-resolved temporal speckle contrast is computed from the variance-to-mean-squared ratio of the TOF-gated interferometric signal after integration over an exposure window *T*. This approach is conceptually closest to conventional speckle-contrast analysis and would be natural in a future parallel implementation, where independent speckle realizations are available across multiple detector pixels or channels.

The second is an indirect estimator based on the TOF-gated field autocorrelation. In this approach, *g*_1_(*τ*_*s*_, *τ*_*d*_) is first computed from repeated short-exposure swept-source measurements and then converted into *κ*^2^(*τ*_*s*_, *T*) using Eq. (9). In the absence of additive noise, the direct and *g*_1_-based estimators yield equivalent temporal speckle-contrast values. In the presence of additive detector noise, however, the direct variance-based estimator is biased because the noise variance contributes directly to the measured contrast. By contrast, the *g*_1_-based estimator localizes the additive-noise contribution primarily at zero lag, allowing the physiological decorrelation component to be recovered from nonzero correlation lags. For this reason, the *g*_1_-based estimator was used for all phantom and *in vivo* analyses in this work, while the direct estimator was retained as a numerical comparator. The difference between the two estimators is quantified in Section 4.2.

## 3. Methods

### 3.1. Numerical simulations

DTOF curves were generated for a semi-infinite homogeneous medium using the diffusion-theory expression of Patterson et al. [28], with absorption *µa*, reduced scattering *µ*_*s*_′, refractive index n = 1.4, and a 10 mm source–detector separation. Three scattering values (*µ*_*s*_′ = 7.5, 10, 12.5 *cm*^−1^) bracketed the range of scattering properties.

TOF-resolved sample fields *U*_*s*_(*r, τ*_*s*_, *t*_*d*_) were synthesized as sequences of correlated speckle frames using the Duncan–Kirkpatrick algorithm [29], with each TOF bin assigned a decorrelation rate *ξ*(*τ*_*s*_) prescribed by Eq. (11) (homogeneous medium) or by the layer-dependent *µ*_*s*_′ through Eq. (11) in the bi-layer medium. A uniform reference field *U*_*r*_(*r, t*_*d*_) of amplitude *a*_*r*_ ≫ |*U*_*s*_| was added coherently, together with an additive noise field *U*_*r*_ (*r, t*_*d*_) with prescribed amplitude *a*_*n*_ ∈ {0, 0.1, 0.25, 0.5}. Synthetic interferometric intensity samples were obtained as |*U*_*s*_ + *a*_*r*_ *U*_*r*_ + *a*_*n*_ *U*_*n*_|^2^ and Fourier-transformed along *v* to recover the TOF axis. Speckle contrast was computed with both estimators (Section 2.4); the analytical *κ*^2^ (Eq. 9) served as ground truth.

Bi-layer simulations were then performed to reproduce the layered phantom configuration, in which a superficial solid scattering slab was placed over a deeper liquid layer. The first layer had thickness *d* ∈ {10,15} mm and reduced scattering coefficient 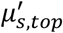, while the second layer had reduced scattering coefficient 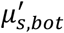. The two simulated scattering configurations matched the phantom cases used experimentally: 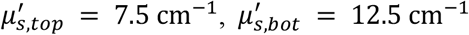, and the reversed case 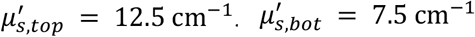. As described by Eq. (11), the TOF-dependent decorrelation rate scales with *µ*_*s*_′, *αD*_*B*_, and photon time of flight. Therefore, changing the top and bottom reduced scattering coefficients directly changed the predicted decorrelation behaviour of photons sampling each layer. This configuration was used to test whether late TOF gates become preferentially sensitive to the deeper layer and whether the transition predicted by the DTOF/path-length distribution could be recovered experimentally.

### 3.2. Optical setup

The experimental system was implemented as a single-channel interferometric near-infrared spectroscopy (iNIRS) Mach–Zehnder interferometer (Fig. 2A). The light source was a tunable laser diode centered at 780 nm (EYP-DFB-0780-00020-1500-BFY12-0005, Eagleyard-Toptica) with a linewidth of 0.6 MHz, corresponding to FWHM ≈ 1.2 fm. The laser output was divided into sample and reference arms using a 90:10 fiber coupler. The sample beam illuminated the target tissue through a single-mode fiber (the optical power at the fiber output was maintained below 15 mW), and the diffusely backscattered light was collected into a single-mode fiber. The reference beam propagated in a single-mode fiber and was combined with the measurement beam in a 2×2 50:50 optical coupler. The two coupler outputs were connected to a balanced photodetector (PDB450A-AC, Thorlabs with the 45 MHz bandwidth), which converted the optical interference signal into an electrical voltage. The output signal was digitized using an acquisition card (M2p.5962-x4, Spectrum Instrumentation, configured for 100 MS/s at 14-bit depth) connected to a PC, where it was processed numerically using a GPU-accelerated code. For all measurements we used a sweep rate of 50 kHz, resulting in 2000 spectral points per sweep.

**Figure 2.**
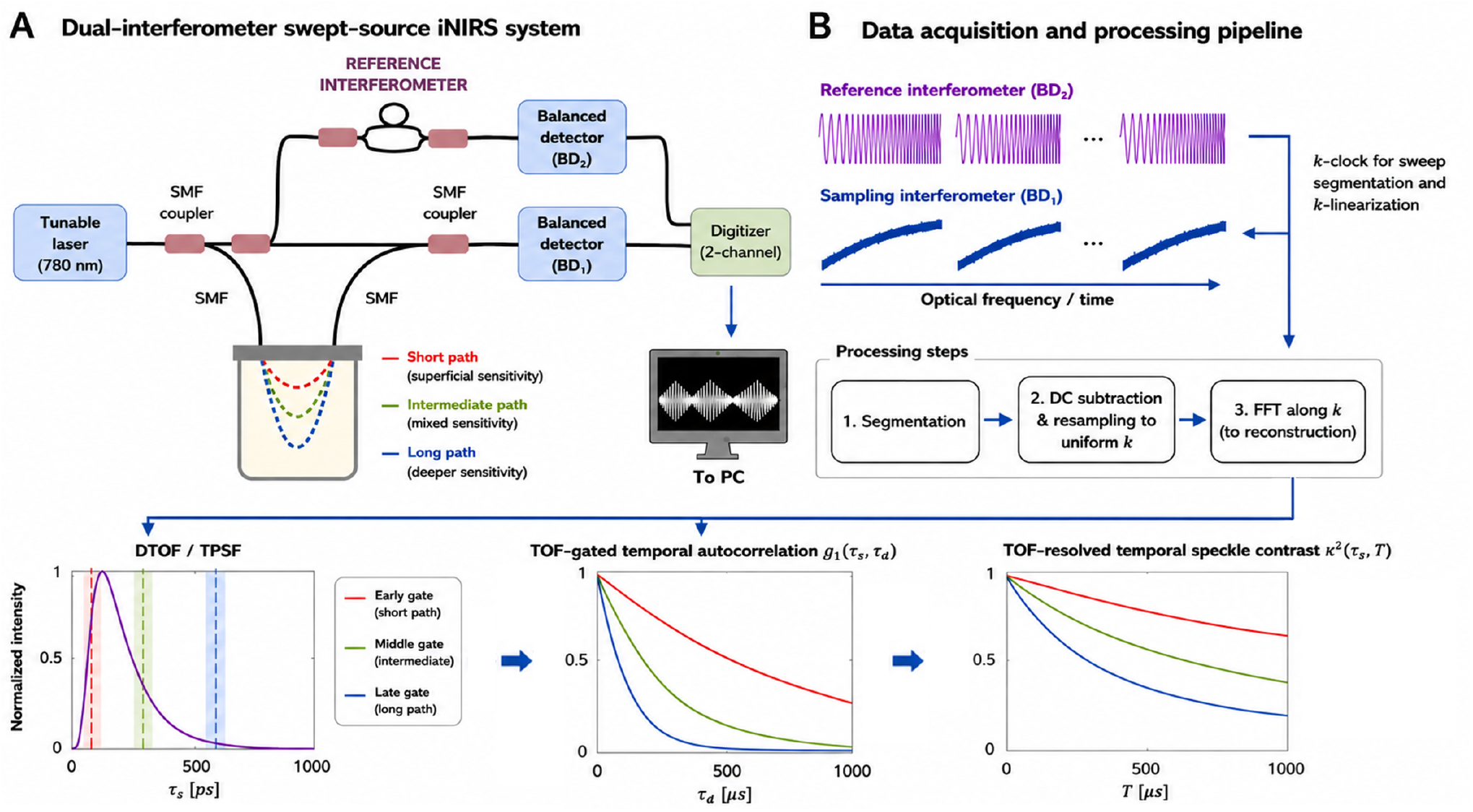
Experimental setup and acquisition pipeline. (A) Dual-interferometer swept-source iNIRS architecture. A fibre-coupled tunable distributed-feedback laser diode operating near 780 nm is first split into sample and reference paths of the measurement interferometer. The sample arm illuminates the target through a single-mode fibre, and diffusely backscattered light is collected into another single-mode fibre. The collected sample field is recombined with the measurement-reference field in a 2 × 2, 50:50 fibre coupler and detected by balanced photodetector BD_1_. A small fraction of the reference path is tapped off and sent to a separate reference interferometer with a fixed optical path-length difference, whose outputs are detected by balanced photodetector BD_2_. The BD_2_ trace provides the k-clock signal used for sweep segmentation and wavenumber linearization. Both balanced-detector signals are digitized simultaneously by a two-channel acquisition card and processed offline. (B) Acquisition and processing pipeline. The reference-interferometer phase is used to segment individual sweeps, correct laser-sweep nonlinearity, and resample the BD_1_ measurement interferogram onto a uniform wavenumber grid. After DC subtraction and Fourier transformation along k, the TOF-resolved complex field and the corresponding DTOF/TPSF are recovered. TOF gates are selected from the DTOF, and the temporal field autocorrelation *g*_1_(*τ*_*s*_, *τ*_*d*_) is computed within each gate across consecutive sweeps. TOF-resolved temporal speckle contrast, *κ*^2^(*τ*_*s*_, *T*), is then estimated indirectly from *g*_1_ using the speckle-visibility relation.

A fibre-based reference interferometer was included to provide the wavenumber-calibration signal. This interferometer had a fixed optical path-length difference of 1 m and generated a periodic interference signal during each laser sweep. The reference-interferometer trace served two purposes. First, it enabled segmentation of individual sweeps from the measurement interferometer. Second, its phase was used to correct laser-sweep nonlinearity and to resample the iNIRS interferometric trace onto a uniform wavenumber grid prior to Fourier transformation. This ensured accurate reconstruction of the TOF-resolved complex field and the corresponding DTOF/TPSF [4,26]. Because the present implementation used a single detection channel, the experiments did not provide ROI-based spatial speckle ensembles or thousands of independent camera-pixel channels. Therefore, the reported TOF-resolved temporal speckle contrast was obtained from the *g*_1_-based analysis of repeated single-channel iNIRS measurements.

### 3.3. Liquid and bi-layer phantoms

Homogeneous liquid phantoms were prepared by diluting homogenized milk (3.2% fat) in distilled water to two reduced-scattering values (*µ*_*s*_′ = 7.5, 12.5 cm^− 1^), with a constant absorption *µ*_*a*_ = 0.03 cm^−1^. These values matched the scattering contrasts used in the bi-layer phantom simulations. Optical properties were validated independently with a TD-NIRS reference system [30]. The phantom was held in a custom, 3D-printed, semi-infinite cubic container, and the source and detector fibres were positioned in a 3D-printed holder at a source–detector separation of *ρ* = 1.0 cm.

To experimentally validate the proposed TOF-resolved analysis, we prepared bi-layer phantoms with opposing scattering contrasts. In Case A, the top layer had lower reduced scattering than the bottom layer 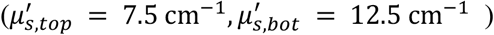, whereas in Case B the contrast was reversed 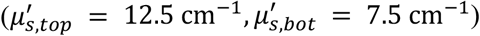. The top-layer thickness was d = 10 and 15 mm for both cases.

### 3.4. In vivo measurements

Two proof-of-concept in vivo paradigms were performed. First, forearm blood-flow modulation was induced using a cuff-occlusion protocol in a 39-year-old female volunteer, Fitzpatrick skin type I. The optode was secured on the volar side of the left forearm with an elastic strap. A 2 min baseline was followed by 2 min of suprasystolic occlusion at 180 mmHg and a 2 min post-release period to capture reactive hyperemia. Second, prefrontal activation was assessed during a Sudoku task in a 33-year-old male volunteer, Fitzpatrick skin type I. The probe was placed on the left forehead, and the volunteer alternated between rest, Sudoku, and post-task rest periods, each lasting 2 min. In both paradigms, the source–detector separation was 10 mm.

All procedures were approved by the Commission of Bioethics at the Military Institute of Medicine, Poland, permission no. 90/WIM/2018, and were conducted in accordance with the Declaration of Helsinki. Written informed consent was obtained before each measurement session.

### 3.5. Data processing

Offline processing followed the TOF-iSCOS framework described in Section 2. First, the DC offset was removed from the single-channel interferometric trace by moving-average subtraction along the optical-frequency sweep. Second, the reference-interferometer phase was used to correct sweep nonlinearity and resample the measurement interferogram onto a uniform wavenumber grid. Third, an inverse Fourier transform along the optical-frequency axis was applied to each sweep, yielding the TOF-resolved complex interferometric field *S*(*τ*_*s*_, *t*_*d*_), where *t*_*d*_ indexes consecutive swept-source acquisitions.

The distribution of times of flight was estimated from the time-averaged TOF-resolved intensity,

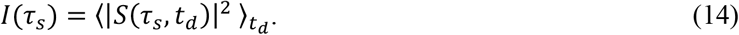

TOF gates were then selected from the DTOF to represent early, intermediate, and late photon arrival times. Within each gate, the temporal field autocorrelation was computed from the TOF-resolved complex field as

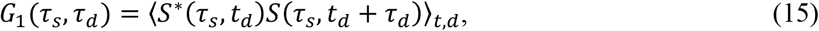

and normalized by the zero-lag value to obtain

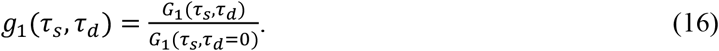

Finally, TOF-resolved temporal speckle contrast, *κ*^2^(*τ*_*s*_, *T*), was estimated indirectly from *g*_1_(*τ*_*s*_, *τ*_*d*_) using the speckle-visibility relation in Eq. (9), with synthetic exposure times *T* = *N*Δ*t* constructed from consecutive swept-source acquisitions. The direct variance-based estimator was used only in numerical simulations as a comparator and was not used for the main phantom or *in vivo* analyses.

Fits used a two-parameter exponential plus offset for g_1_,

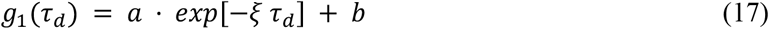

and the corresponding speckle-contrast model

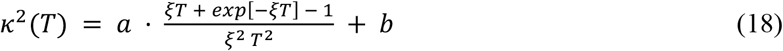

To estimate the dynamic coefficient, *αD*_*B*_, the measured *g*_1_(*τ*_*s*_, *τ*_*d*_) curves were fitted with the single-layer DWS prediction using adaptive fitting windows selected by RMSE minimization, as in CW-iSCOS [24]. The TOF-resolved *αD*_*B*_ was obtained by inverting Eq. (11), 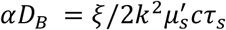, where *k* is the in-medium wavenumber and c is the speed of light in the medium. In simulations and phantom analyses, *αD*_*B*_ is interpreted as a dynamic coefficient. In the in vivo forearm and forehead measurements, the same recovered quantity is interpreted as a blood-flow index (BFI), following conventional DWS/DCS terminology. In simulations and phantoms, 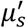 was set by the nominal or independently measured optical properties. *In vivo*, 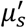 was fitted from each baseline TPSF using the Patterson semi-infinite diffusion model [28] convolved with the measured instrument response function. Relative *in vivo* changes were computed as ΔBFI/BFI_0_, with BFI_0_ averaged over the corresponding baseline window.

## 4. Results

### 4.1. Simulation validation: TOF-resolved *κ*^2^(***τ***_***s***_, ***T***)

Figure 3 illustrates the TOF-iSCOS forward model for three reduced scattering coefficients 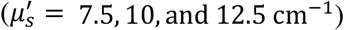. The left column shows the simulated diffuse time-of-flight distributions (DTOFs), together with three representative TOF gates (*τ*_*s*_ = 150, 400, and 650 ps). As expected, increasing *µ*_*s*_′ broadens the DTOF and shifts photon probability toward longer path lengths, reflecting stronger multiple scattering within the medium.

**Figure 3.**
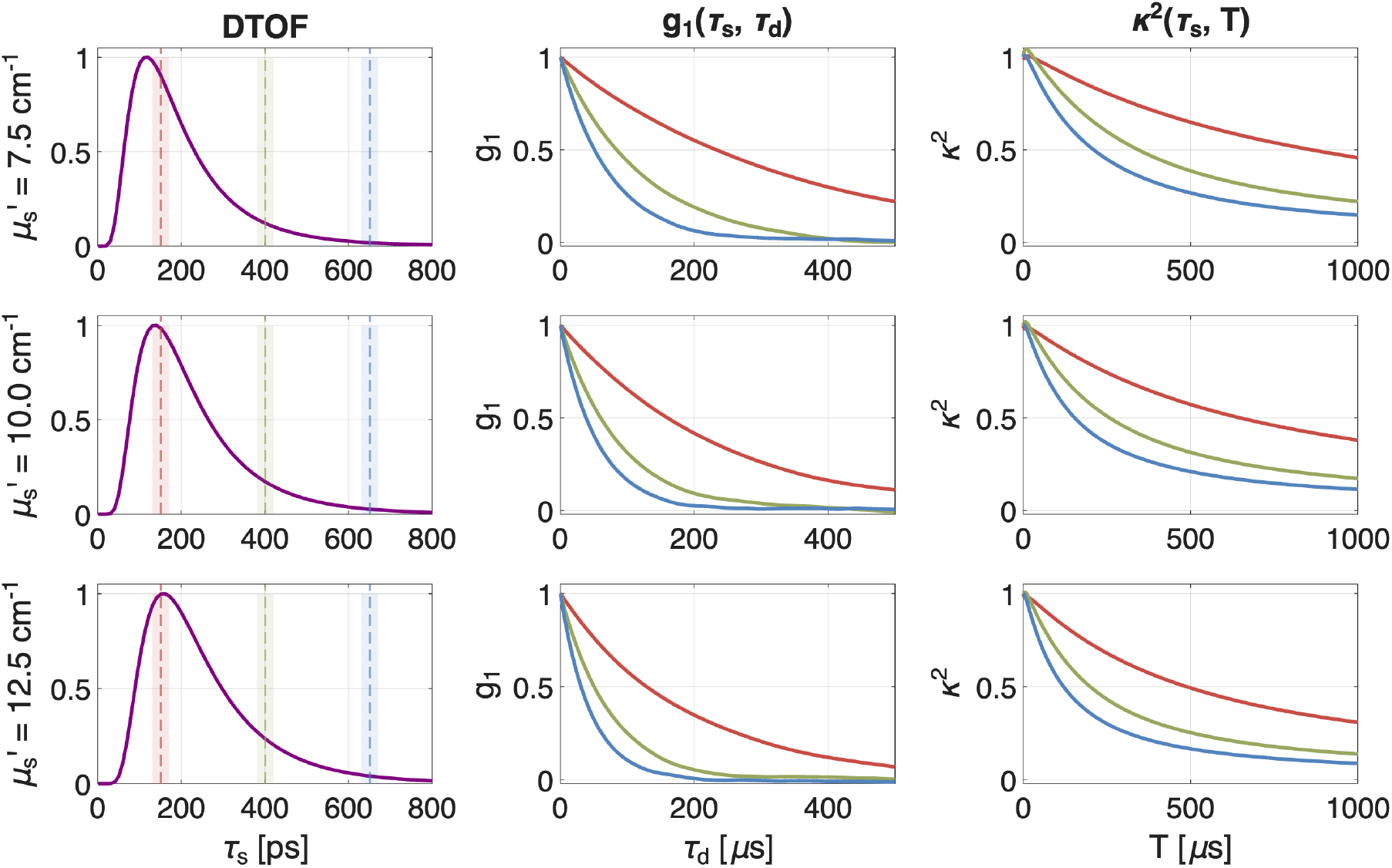
Simulated TOF-iSCOS forward model in a homogeneous semi-infinite medium for three reduced scattering coefficients (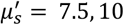, 00, and 12.5 cm^−1^). Rows correspond to different scattering coefficients. Left column: normalized diffuse time-of-flight distributions (DTOFs) with three representative TOF gates (*τ*_*s*_ = 050, 400, and 650 ps) indicated by dashed vertical lines. Middle column: TOF-resolved field autocorrelation functions *g*_1_(*τ*_*s*_, *τ*_*d*_) at the selected gates. Right column: corresponding TOF-resolved speckle-contrast curves *κ*^2^(*τ*_*s*_, *T*). For all scattering coefficients, later TOF gates decorrelate more rapidly and exhibit faster contrast decay.

The middle column shows the corresponding TOF-resolved field autocorrelation functions *g*_1_(*τ*_*s*_, *τ*_*d*_). For all scattering coefficients, later TOF gates exhibit faster decorrelation than earlier gates. This behavior follows directly from the TOF dependence of the effective photon path length: photons detected at longer TOFs experience a larger number of dynamic scattering events and therefore accumulate greater phase fluctuations. Consequently, the decorrelation rate increases monotonically with TOF.

The right column presents the corresponding TOF-resolved speckle-contrast curves *κ*^2^(*τ*_*s*_, *T*) obtained from Eq. (9). The ordering observed in *g*_1_(*τ*_*s*_, *τ*_*d*_) is preserved in the exposure-time domain, with late TOF gates exhibiting the fastest contrast decay and early TOF gates the slowest. Importantly, this ordering is maintained across all simulated scattering coefficients, demonstrating that TOF gating provides a robust mechanism for separating photon populations with different dynamical sensitivities. These simulations validate the expected relationship between photon time-of-flight, field decorrelation, and exposure-dependent speckle visibility that underpins the TOF-iSCOS framework.

### 4.2. Simulation: noise robustness of the two estimators

Figure 4 compares the direct variance-based estimator and the *g*_1_-based estimator of TOF-resolved temporal speckle contrast, *κ*^2^(*τ*_*s*_, *T*), under additive detector noise. Simulations were performed for three TOF gates and three additive-noise amplitudes. In the absence of additive noise, both estimators recovered the theoretical exposure dependence, confirming consistency between the simulated speckle dynamics, the analytical *κ*^2^ model, and the numerical estimators.

**Figure 4.**
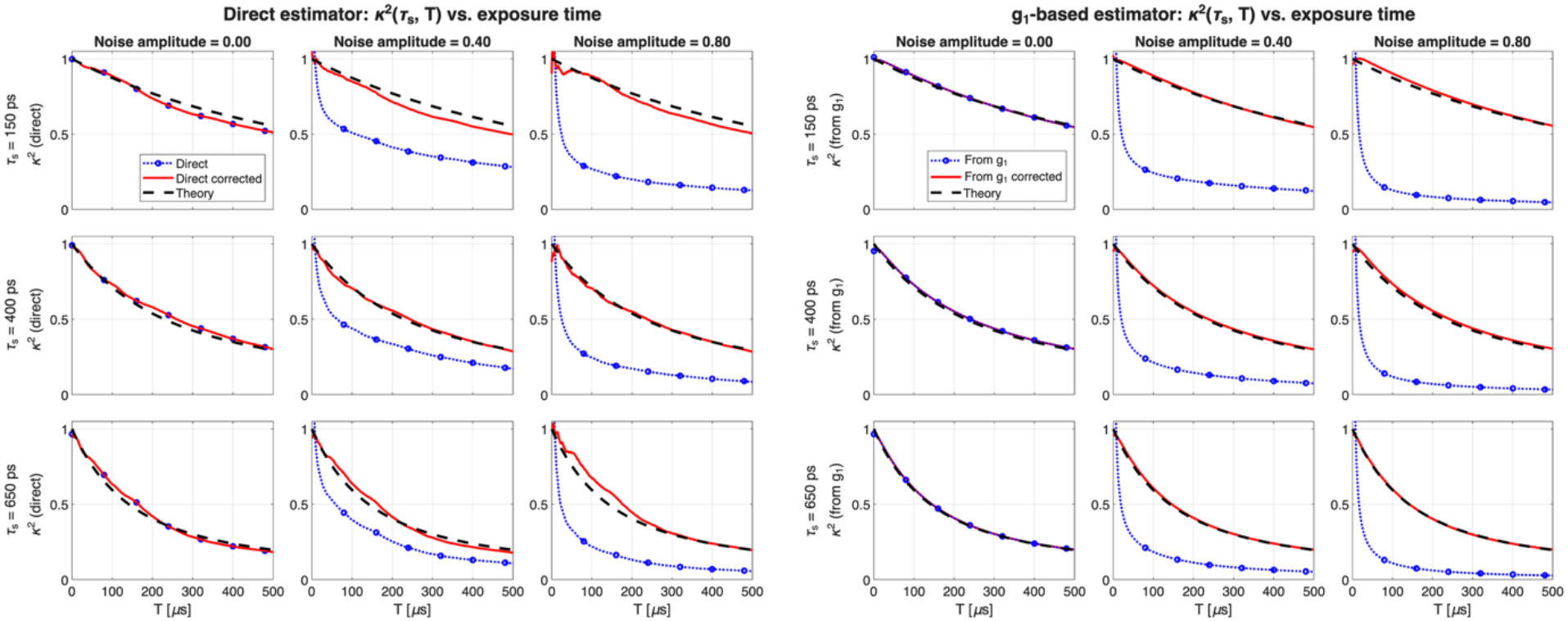
Noise robustness of the direct and *g*_1_-based estimators of *κ*^2^(*τ*_*s*_, *T*). Simulations are shown for three TOF gates and three additive-noise amplitudes. In the direct estimator, additive noise biases the variance-based *κ*^2^ estimate, and standard variance subtraction reduces but does not fully eliminate the bias under limited speckle statistics. In the *g*_1_-based estimator, additive noise is corrected at the correlation level by reconstructing the zero-lag autocorrelation value from nonzero lag points, preserving agreement with theory across noise levels.

With increasing additive noise, the uncorrected direct estimator became biased because the noise contribution entered the variance used to estimate *κ*^2^. Standard variance subtraction, analogous to shot- and dark-noise correction used in SCOS, partially compensated this effect. However, its performance depended on the number of statistically independent speckle realizations and on accurate knowledge of the noise variance. Residual finite-sample variance and imperfect noise subtraction therefore propagated into the final contrast estimate.

By contrast, the *g*_1_-based estimator remained close to theory after correlation-level correction. In this approach, the contribution from additive uncorrelated noise is concentrated predominantly at zero correlation lag. Reconstructing the zero-lag autocorrelation value by extrapolation from nonzero lag points, *τ*_*d*_ > 0, therefore removes the dominant noise contribution before evaluating the exposure-time integral in Eq. (9). The advantage of the *g*_1_-based estimator was most pronounced at the highest noise amplitude and in TOF gates where the dynamic speckle contribution was weakest relative to the noise floor.

To evaluate the effect of detector parallelism, the simulations were repeated for synthetic detector arrays ranging from 32 × 32 to 256 × 256 pixels (Fig. S1). The error of the direct estimator decreased substantially with increasing detector count, confirming that finite speckle statistics are a major contributor to the residual error observed in Fig. 4. Standard variance subtraction further improved performance, but the direct estimator remained more sensitive to detector size than the *g*_1_-based approach. By contrast, the corrected *g*_1_-based estimator maintained low error across all simulated detector sizes, indicating that its robustness originates primarily from correlation-level noise correction rather than detector averaging.

These results suggest a practical division between the two estimators. In future highly parallel implementations, where many independent speckle realizations are available, the direct estimator can converge toward the theoretical *κ*^2^(*τ*_*s*_, *T*) curve after appropriate variance correction. In single-detector or low-channel-count swept-source iNIRS implementations, the *g*_1_-based estimator is preferable because additive-noise correction is performed at the autocorrelation level before *κ*^2^ is computed.

### 4.3. Simulation: bi-layer geometry and *ξ*(*τ*_*s*_) recovery

Figure 5 illustrates the expected TOF-iSCOS response in a bi-layer medium consisting of a superficial layer with reduced scattering coefficient 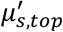 above a deeper layer with reduced scattering coefficient 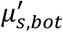 Two representative configurations were simulated. Case A: 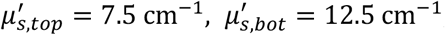, case 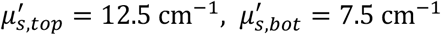. For both cases, superficial layer thicknesses of d = 10 mm and d = 15 mm were simulated.

**Figure 5.**
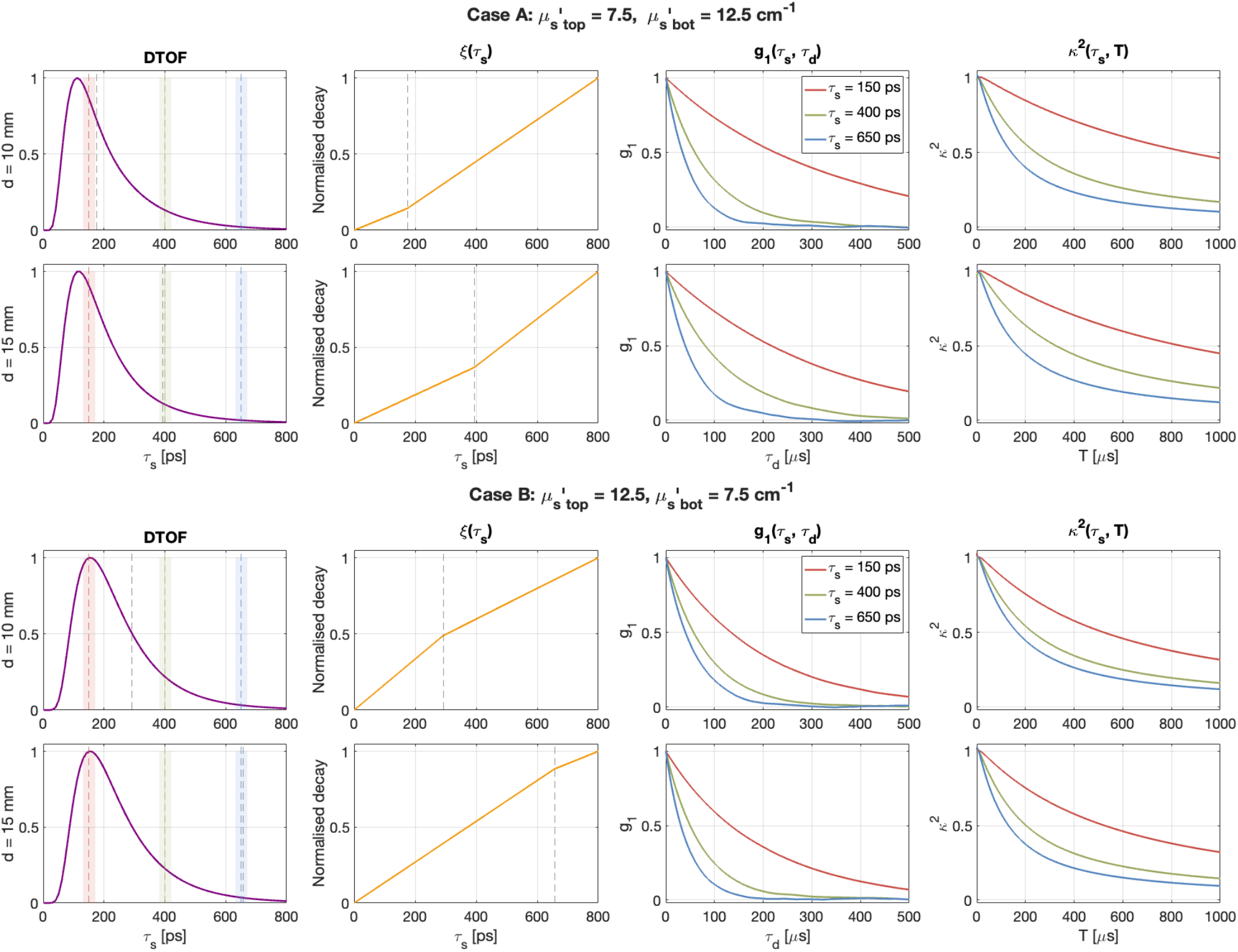
Bi-layer TOF-iSCOS simulations. Top panels: Case A 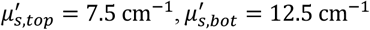. Bottom panels: Case B 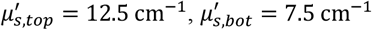. For each case, superficial-layer thicknesses of d = 10 and 15 mm are shown. Columns show the DTOF, TOF-dependent decorrelation rate *ξ*(*τ*_*s*_), TOF-resolved field autocorrelation *g*_1_(*τ*_*s*_, *τ*_*d*_), and TOF-resolved temporal speckle contrast *κ*^2^(*τ*_*s*_, *T*). Vertical dashed lines indicate representative TOF gates. The onset and slope transition in *ξ*(*τ*_*s*_) arise from the layer-dependent reduced scattering coefficient in Eq. (11), marking the TOF range over which photons begin to sample the deeper layer and producing gate-dependent changes in both *g*_1_ and *κ*^2^.

The DTOFs show relatively subtle differences between the two configurations and provide only indirect evidence of the layer transition. In contrast, the TOF-dependent decorrelation rate, *ξ*(*τ*_*s*_), shows a clear change in slope at photon times of flight where the detected paths begin to sample the second layer substantially. This behaviour follows directly from Eq. (11): for a given dynamic coefficient, *ξ*(*τ*_*s*_) scales with 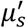 and photon time of flight. Therefore, as late-arriving photons accumulate a larger fraction of their path length in a layer with a different 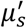 the effective TOF-dependent decorrelation rate changes.

The onset of this transition provides an additional depth-sensitive marker. Because each TOF gate contains an ensemble of photon trajectories, the transition is gradual rather than abrupt. Its onset indicates the earliest TOF range in which a non-negligible fraction of detected photons begins to sample the second layer, while the subsequent change in slope reflects increasing weighting of those deeper paths. The onset shifts toward later TOFs as the superficial layer thickness increases, consistent with the longer path lengths required to reach the deeper region.

The TOF-resolved field autocorrelation functions, *g*_1_(*τ*_*s*_, *τ*_*d*_), and the corresponding temporal speckle-contrast curves, *κ*^2^(*τ*_*s*_, *T*), inherit this TOF-dependent change in decorrelation rate. Early TOF gates are dominated by paths confined mainly to the superficial layer, whereas late gates increasingly include photons that have sampled the deeper layer. Consequently, the separation between gate-specific *g*_1_ and *κ*^2^ curves provides a readout of depth-dependent scattering and decorrelation behavior. Importantly, this separation is observed even when the DTOF itself changes only weakly, indicating that decorrelation-derived quantities can reveal layer-dependent transport effects that are not obvious from photon-arrival distributions alone.

The two bi-layer configurations produce distinct onset and transition behaviors. In Case A, where the superficial layer has the lower reduced scattering coefficient, photons reach the higher-scattering deeper layer at relatively moderate TOFs. As a result, the transition onset appears earlier and the slope of *ξ*(*τ*_*s*_) increases after deeper-layer sampling becomes appreciable. In Case B, the more strongly scattering superficial layer delays photon transport into the lower-scattering deeper region, shifting the transition onset toward later TOFs. In this case, the slope of *ξ*(*τ*_*s*_) decreases after the transition. For the d = 15 mm Case B configuration, the onset occurs mainly in the late tail of the DTOF, showing that sensitivity to the deeper layer can emerge only in a small late-arriving photon population.

These simulations demonstrate that the TOF at which layer sensitivity emerges depends not only on the physical layer thickness but also on the optical properties of the superficial layer. Thus, TOF-resolved decorrelation and temporal speckle-contrast measurements provide access to depth-dependent scattering structure that cannot be inferred reliably from the DTOF alone. This motivates the combination of TOF gating with interferometric decorrelation analysis in TOF-iSCOS.

### 4.4. Simulation: recovery of *ξ*(*τ*_*s*_) and dynamic coefficients

Figure 6 evaluates the ability of TOF-iSCOS to recover the TOF-dependent decorrelation rate, *ξ*(*τ*_*s*_), and the corresponding dynamic coefficient, *αD*_*B*_, from simulated bi-layer measurements. The same bi-layer geometries as in Fig. 5 were used, including Cases A and B and superficial-layer thicknesses of 10 and 15 mm.

**Figure 6.**
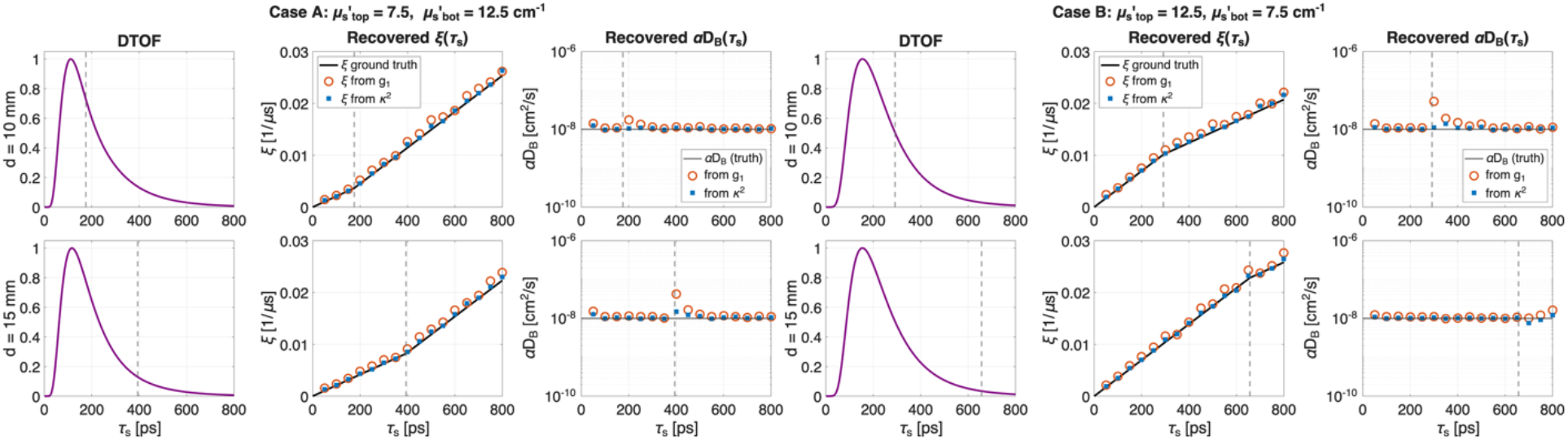
Recovery of the TOF-dependent decorrelation rate *ξ*(*τ*_*s*_) and dynamic coefficient *αD*_*B*_ in simulated bi-layer media. Results are shown for Case A, 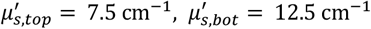, and Case B, 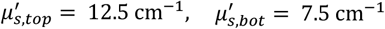. Rows show superficial-layer thicknesses of 10 and 15 mm. The first column shows the normalized DTOF with vertical dashed lines marking the TOF onset. The second column compares the ground-truth decorrelation rate *ξ*(*τ*_*s*_), solid black line, with values recovered from TOF-resolved field autocorrelation analysis, orange circles, and TOF-resolved temporal speckle-contrast analysis, blue squares. The third column shows the corresponding recovered *αD*_*B*_. Both *g*_1_- and *κ*^2^-based recovery identify the layer-dependent transition in *ξ*(*τ*_*s*_). After accounting for the *µ*_*s*_′-dependence of Eq. (11), the recovered *αD*_*B*_ remains close to the imposed dynamic coefficient across the usable TOF range.

The recovered decorrelation rates obtained from TOF-resolved field autocorrelation analysis, *g*_1_(*τ*_*s*_, *τ*_*d*_), and from TOF-resolved temporal speckle-contrast analysis, *κ*^2^(*τ*_*s*_, *T*), are compared with the ground-truth *ξ*(*τ*_*s*_) used in the forward model. In all cases, both approaches closely follow the ground truth and correctly identify the onset and location of the layer-dependent transition in *ξ*(*τ*_*s*_). Increasing the superficial-layer thickness shifts the transition toward longer TOFs, consistent with the longer photon path lengths required before the detected trajectories substantially sample the deeper layer.

The *κ*^2^-based recovery is particularly important for TOF-iSCOS because it demonstrates that the exposure-dependent temporal speckle-contrast curves preserve sufficient information to recover the TOF-dependent decorrelation rate. The agreement between *ξ*(*τ*_*s*_) recovered from *g*_1_ and from *κ*^2^ confirms the consistency of the two analysis routes described by the speckle-visibility relation. Minor deviations occur mainly near the transition region and at the latest TOFs, where the DTOF intensity is lowest and the effective contribution of late photon paths is more weakly sampled.

The recovered *αD*_*B*_ profiles provide a consistency check on the quantitative interpretation of the decorrelation rate. Because the simulated layers differ in reduced scattering coefficient rather than in the imposed dynamic coefficient, the transition in *ξ*(*τ*_*s*_) arises from the 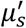-dependence in Eq. (11). After accounting for this scattering dependence, the recovered *αD*_*B*_ remains close to the imposed value across the usable TOF range for both *g*_1_- and *κ*^2^-based recovery. These results demonstrate that TOF-iSCOS can recover both the TOF-dependent decorrelation structure and the underlying dynamic coefficient when the scattering-dependent contribution to *ξ*(*τ*_*s*_) is properly accounted for.

### 4.5. Homogeneous liquid phantoms

Figure 7 presents experimental TOF-resolved measurements acquired in homogeneous liquid phantoms with reduced scattering coefficients 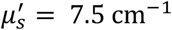 and 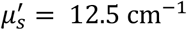 at a source–detector separation of 10 mm. The measured DTOFs show the expected broad temporal distribution of photon arrival times and provide sufficient dynamic range to define representative early, middle, and late TOF gates.

**Figure 7.**
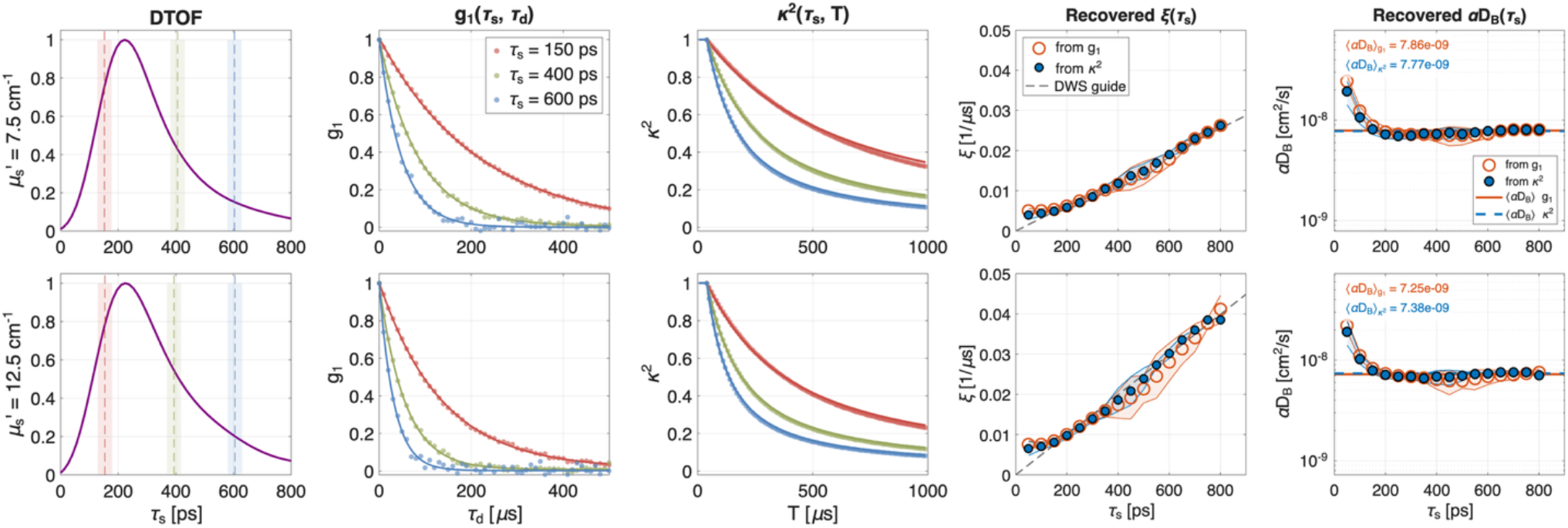
Experimental TOF-resolved measurements in homogeneous liquid phantoms. Results are shown for reduced scattering coefficients 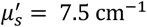 (first row) and 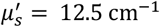 (second row) at a source–detector separation of 10 mm. Columns show the measured DTOF, TOF-gated field autocorrelation functions *g*_1_(*τ*_*s*_, *τ*_*d*_), TOF-resolved temporal speckle-contrast curves *κ*^2^(*τ*_*s*_, *T*), recovered decorrelation rates *ξ*(*τ*_*s*_), and corresponding recovered dynamic coefficients *αD*_*B*_ (*τ*_*s*_). Vertical dashed lines indicate representative early, middle, and late TOF gates. Error envelopes in the recovered *ξ*(*τ*_*s*_) and *αD*_*B*_ (*τ*_*s*_) plots indicate 3 × SD_robust_ with SD_robust_ estimated from MAD. Later TOF gates exhibit faster field decorrelation and stronger temporal contrast decay, consistent with increased photon path lengths. The decorrelation rates recovered from *g*_1_ and from *κ*^2^ agree closely and follow the DWS-guided TOF dependence. After accounting for the *µ*_*s*_′-dependence of Eq. (11), the recovered *αD*_*B*_ remains approximately constant across the usable TOF range, as expected for homogeneous liquid phantoms.

The corresponding TOF-resolved field autocorrelation functions, *g*_1_(*τ*_*s*_, *τ*_*d*_), are shown in the second column. For both scattering coefficients, later TOF gates decorrelate more rapidly than earlier gates. This is consistent with the longer photon path lengths sampled by late-arriving photons and the consequently larger accumulated dynamic phase shifts. The experimentally observed gate ordering agrees with the homogeneous-medium prediction: early gates exhibit slower decorrelation, whereas late gates exhibit faster decorrelation.

The third column shows the TOF-resolved temporal speckle-contrast curves, *κ*^2^(*τ*_*s*_, *T*), obtained from the measured *g*_1_(*τ*_*s*_, *τ*_*d*_) using the speckle-visibility relation. As expected from Eq. (9), the ordering observed in the autocorrelation domain is preserved in the exposure-time domain. Early TOF gates show the slowest contrast decay, whereas late TOF gates show the strongest decay. This confirms that the *g*_1_-based estimator provides experimentally stable TOF-resolved temporal speckle contrast from single-channel swept-source iNIRS measurements.

The fourth column shows the decorrelation rates *ξ*(*τ*_*s*_) recovered from both *g*_1_(*τ*_*s*_, *τ*_*d*_) and *κ*^2^(*τ*_*s*_, *T*). Both recovery routes follow the DWS-guided TOF dependence, confirming the consistency between the autocorrelation-based and speckle-contrast-based analyses. The shaded regions indicate uncertainty intervals corresponding to 3 × SD_robust_, where SD_robust_ was estimated from the median absolute deviation (MAD). Across most of the usable TOF range, the *κ*^2^-based recovery exhibits smaller uncertainty than the *g*_1_-based recovery, indicating that fitting the exposure-dependent temporal speckle-contrast curves provides a stable route to estimating *ξ*(*τ*_*s*_). This is particularly important for TOF-iSCOS because it demonstrates that the speckle-contrast representation preserves sufficient information to recover the underlying TOF-dependent decorrelation rate.

The fifth column shows the corresponding recovered *αD*_*B*_(*τ*_*s*_). After accounting for the 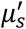 -dependence of Eq. (11), the recovered *αD*_*B*_ remains approximately constant across the usable TOF range for each homogeneous phantom, as expected for a uniform liquid medium. The *κ*^2^-based estimates again show lower variability than the *g*_1_-based estimates over most of the TOF range. Deviations are most apparent at the earliest and latest TOFs, where the DTOF intensity is lower and the estimates are more sensitive to noise, finite sampling, and early- or late-time model limitations. Overall, these measurements validate the TOF-iSCOS processing chain experimentally and confirm that TOF-resolved temporal speckle contrast can robustly recover path-length-dependent dynamics predicted by DWS.

### 4.6. Bi-layer phantoms

Figure 8 tests whether the TOF-dependent decorrelation transitions predicted by the bi-layer simulations can be recovered experimentally. Building on the homogeneous phantom measurements in Fig. 7, which established reference *αD*_*B*_ values for the two liquid scattering conditions, the bi-layer experiments evaluate whether these dynamic coefficients can still be recovered when photons sample a layered scattering structure. Case A represents a lower-scattering superficial layer over a higher-scattering bottom layer, whereas Case B reverses this scattering contrast. For both cases, top-layer thicknesses of d = 10 and 15 mm were evaluated. The DWS-guided curves were generated from Eq. (11) using the nominal layer-specific 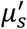 values and the *αD*_*B*_ values independently recovered from the corresponding homogeneous liquid phantoms. Thus, before the estimated transition TOF, the guide follows the top-layer 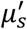, whereas after the transition it follows the bottom-layer 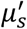.

**Figure 8.**
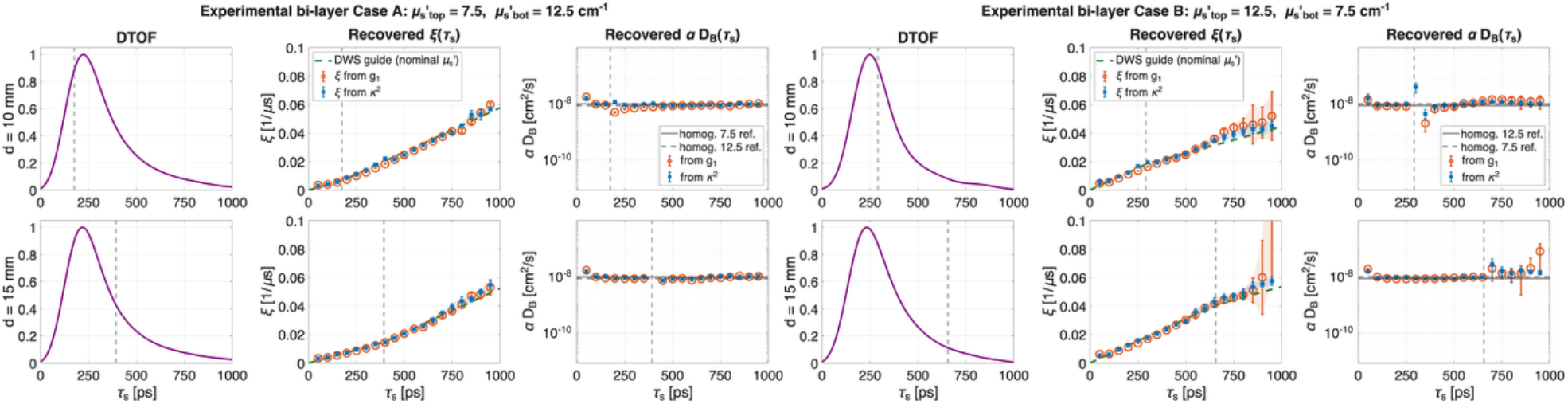
Experimental recovery of TOF-dependent decorrelation trends in bi-layer phantoms. Case A corresponds to 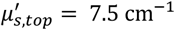 and 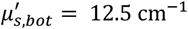, Case B corresponds to the reversed scattering contrast, 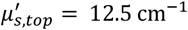 and 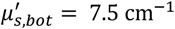. Rows show top-layer thicknesses of d = 10 and 15 mm. Columns show the measured DTOF, recovered TOF-dependent decorrelation rate *ξ*(*τ*_*s*_), and recovered dynamic coefficient *αD*_*B*_ (*τ*_*s*_). Dashed vertical lines indicate the estimated transition TOF. Green dashed curves show DWS-guided trends generated from Eq. (11) using the nominal layer-specific 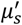 values and the *αD*_*B*_ values independently recovered from the corresponding homogeneous liquid phantoms. Horizontal reference lines in the *αD*_*B*_ panels indicate the homogeneous-phantom *αD*_*B*_ values established in Fig. 7, which serve as independent reference values for the bi-layer recovery. Error bars and shaded envelopes indicate 3 × SD_robust_, with SD_robust_ estimated from the median absolute deviation (MAD). Both *g*_1_- and *κ*^2^-derived recovery reproduce the expected direction of the TOF-dependent transition: increased post-onset *ξ*(*τ*_*s*_) slope in Case A and reduced or flattened post-onset slope in Case B. The *κ*^2^-based recovery generally shows lower uncertainty, particularly in late TOF gates.

In Case A, the recovered *ξ*(*τ*_*s*_) follows the expected increase in slope as late-arriving photons progressively sample the more strongly scattering lower layer. The transition is visible for both top-layer thicknesses, with the onset shifted toward later TOFs for the thicker superficial layer. For d = 10 mm, the transition onset occurs at relatively early TOFs, indicating that at a 10 mm source–detector separation a substantial fraction of the detected photons already samples the lower layer. Both *g*_1_- and *κ*^2^-derived estimates reproduce the DWS-guided trend, and the recovered *αD*_*B*_ (*τ*_*s*_) remains close to the homogeneous-phantom reference values after accounting for the layer-dependent 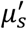.

In Case B, the reversed scattering contrast produces the opposite transition behavior. For the 10 mm top layer, the recovered *ξ*(*τ*_*s*_) initially follows the high-scattering superficial-layer trend and then flattens relative to that trend after the estimated transition TOF, consistent with increasing photon sampling of the lower-scattering bottom layer. For the 15 mm top layer, the estimated transition occurs in the late DTOF tail, leaving fewer high-SNR post-onset points. In this regime, the *κ*^2^-derived recovery remains more consistent with the predicted delayed transition, whereas the *g*_1_-based recovery becomes unstable at the latest analyzed TOFs, producing large uncertainty and an apparent excursion in the recovered parameters.

The recovered *αD*_*B*_(*τ*_*s*_) provides an additional consistency check against the homogeneous-reference measurements. Because the bi-layer phantoms differ primarily in 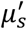, the expected transition should appear mainly in *ξ*(*τ*_*s*_), whereas the recovered *αD*_*B*_ should remain close to the dynamic coefficients measured independently in the homogeneous liquid phantoms. This behavior is observed across most of the usable TOF range, indicating that the TOF-iSCOS analysis can recover the expected dynamic coefficient even when the detected photon paths traverse a layered scattering structure. Deviations occur near the estimated transition and in the late DTOF tail, where signal levels are lower and the recovered parameters are more sensitive to noise and finite sampling.

As in the homogeneous phantom measurements, the *κ*^2^-based recovery generally shows lower uncertainty than the *g*_1_-based recovery. This advantage is particularly clear in late TOF gates, where the DTOF intensity is low and the field-autocorrelation fit becomes less stable. The temporal speckle-contrast representation therefore provides a more robust experimental route to recovering *ξ*(*τ*_*s*_) in low-SNR TOF regions, supporting its use as the primary TOF-iSCOS readout.

Overall, these measurements demonstrate that TOF-iSCOS can experimentally recover the direction, onset, and relative change of TOF-dependent decorrelation trends in layered scattering media. The transition is more apparent in *ξ*(*τ*_*s*_) than in the DTOF alone, confirming that TOF-resolved decorrelation and temporal speckle-contrast analysis provide enhanced sensitivity to subsurface scattering structure.

### 4.7. Forearm cuff-occlusion challenge

Figure 9 presents an in vivo TOF-iSCOS measurement during a forearm cuff-occlusion challenge. Three representative TOF gates, *τ*_*s*_ = 050, 400, and 650 ps, were selected from the analyzed range 000 ≤ *τ*_*s*_ ≤ 900 ps to illustrate the temporal evolution of the decorrelation rate during baseline, cuff inflation, and cuff release. The primary analysis used the *κ*^2^-based estimator, in which TOF-resolved temporal speckle-contrast curves were obtained from *g*_1_ using the speckle-visibility relation and subsequently fitted to recover *ξ*.

**Figure 9.**
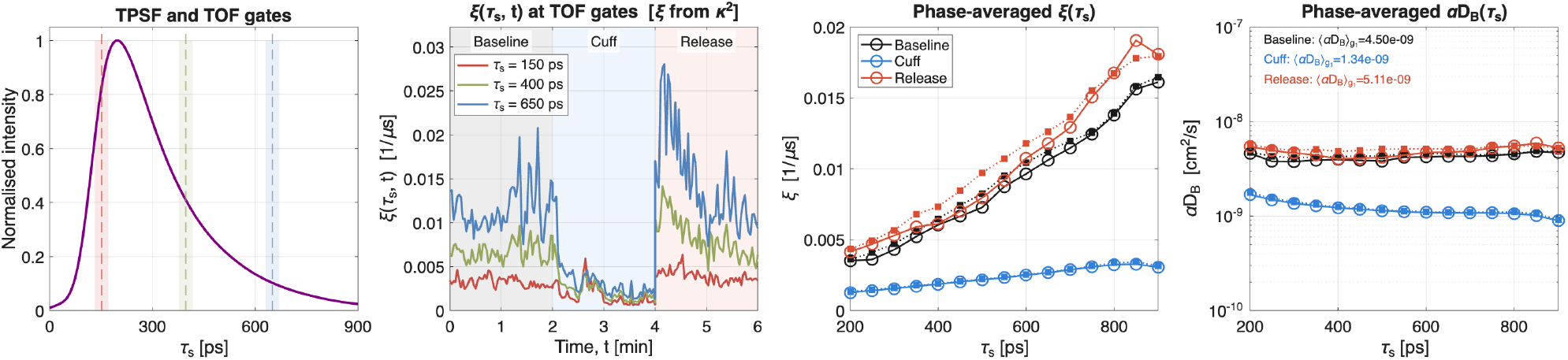
In vivo TOF-iSCOS forearm cuff-occlusion measurement. Columns show, from left to right: the mean TPSF with the analyzed TOF range, and representative TOF gates at *τ*_*s*_ = 050, 400, and 650 ps; the time-resolved decorrelation rate *ξ*(*τ*_*s*_, *t*) recovered with the *κ*^2^-based estimator at the selected TOF gates during baseline, cuff inflation, and cuff release; phase-averaged TOF-dependent decorrelation profiles *ξ*(*τ*_*s*_) for the three physiological phases; and the corresponding phase-averaged blood-flow-index profiles *αD*_*B*_(*τ*_*s*_), computed from Eq. (11) using 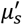 estimated from the baseline TPSF. Shaded bands and error bars indicate robust fitting uncertainty. Supporting fits for the *κ*^2^-based analysis are provided in Fig. S2. The corresponding *g*_1_-based analysis is provided in Fig. S3 and shows the same qualitative response, but with a slightly lower recovered hyperemic peak amplitude.

The time-resolved decorrelation rates *ξ*(*t*) for the selected TOF gates are shown in the second column of Fig. 9. During baseline, *ξ* increases with TOF, with the latest gate exhibiting the largest values. Upon cuff inflation, *ξ* decreases rapidly in all gates and approaches a common low-flow state, consistent with suppression of microvascular perfusion. Following cuff release, all gates exhibit a reactive-hyperemia response, with the largest absolute increase observed at the latest TOF gate. This TOF-dependent amplification is consistent with the greater sensitivity of longer photon trajectories to deeper and more strongly perfused vascular compartments.

The phase-averaged TOF-dependent decorrelation profiles are summarized in the third column of Fig. 9. During baseline and release, *ξ*(*τ*_*s*_) increases monotonically with TOF, whereas during cuff inflation the profile collapses to substantially lower values across the analyzed TOF range. The positive TOF dependence observed during baseline and release mirrors the behavior observed in the phantom measurements, indicating that later-arriving photons accumulate stronger dynamic phase perturbations than earlier-arriving photons.

To convert the recovered decorrelation rates into *αD*_*B*_, Eq. (11) was inverted using 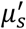 estimated from the baseline TPSF of the forearm measurement rather than nominal phantom values. Specifically, the baseline TPSF was fitted with the Patterson semi-infinite diffusion model convolved with the measured instrument response function, and the resulting 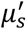 was used in the DWS prefactor for the *αD*_*B*_ profiles.

The corresponding TOF-dependent blood-flow-index profiles are shown in the fourth column of Fig. 9. *αD*_*B*_ decreases during cuff inflation and recovers during release. Baseline and release exhibit a positive TOF dependence, whereas the cuff phase remains suppressed across the analyzed range. These results demonstrate that the *κ*^2^-based TOF-iSCOS estimator resolves the expected physiological modulation during cuff occlusion while preserving the TOF dependence of the recovered flow index.

Supporting fits for the *κ*^2^-based analysis are provided in Fig. S2. The corresponding direct *g*_1_-based analysis is shown in Fig. S3 and reproduces the same qualitative physiological response: suppression during cuff inflation and recovery after cuff release, with stronger modulation in later TOF gates. However, the *g*_1_-based recovery yields a slightly lower hyperemic peak amplitude than the *κ*^2^-based estimator, particularly in the late gate. This difference is consistent with the phantom results, where the *κ*^2^-based recovery provided a more stable estimator in low-SNR TOF regions. We therefore use the *κ*^2^-based recovery as the primary functional readout.

### 4.8. Prefrontal measurement during a Sudoku task

Figure 10 presents a proof-of-concept in vivo TOF-iSCOS measurement acquired from the left forehead during a Sudoku task. The volunteer alternated between three 2 min phases: quiet baseline, Sudoku solving, and recovery. Three TOF gates, *τ*_*s*_ = 400, 600, and 800 ps, were selected to sample progressively later photon populations. This single-subject recording is interpreted as a methodological feasibility demonstration rather than a quantitative cognitive-neuroscience study. Its purpose was to test whether the TOF-dependent functional signature established in simulations, phantoms, and the forearm cuff challenge could also be observed in a forehead activation paradigm.

**Figure 10.**
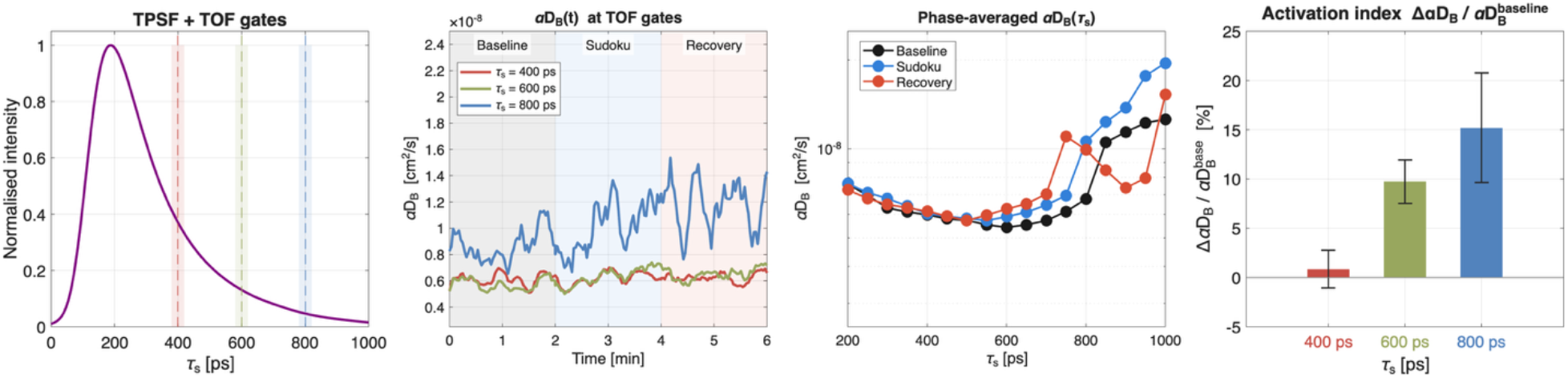
In vivo TOF-iSCOS measurement during a forehead Sudoku task. Columns show, from left to right: the mean TPSF with representative TOF gates at *τ*_*s*_ = 400, 600, and 800 ps; the temporal evolution of *αD*_*B*_ (*t*) recovered with the *κ*^2^-based estimator at the selected TOF gates during baseline, Sudoku, and recovery phases; phase-averaged TOF-dependent *αD*_*B*_ (*τ*_*s*_) profiles for the three phases; and the activation index, 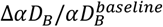, at each TOF gate. Error bars indicate the robust SEM of the Sudoku-period relative changes, computed as 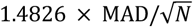, where *N* is the number of valid Sudoku windows. The *κ*^2^-derived response increased with TOF, from +0.8 ± 1.9% at 400 ps to +9.8 ± 2.2% at 600 ps and +15.2 ± 5.6% at 800 ps.

The primary analysis used the *κ*^2^-based estimator. For each TOF gate, *g*_1_(*τ*_*s*_, *τ*_*d*_, *t*) was estimated over 1s temporal windows and converted into TOF-resolved temporal speckle contrast using the speckle-visibility relation. The resulting *κ*^2^(*τ*_*s*_, *T, t*) curves were fitted to recover *ξ*(*τ*_*s*_, *t*), which was then converted into *αD*_*B*_ (*t*) using the optical properties estimated from the baseline TPSF. The supporting fits for the selected TOF gates are shown in Fig. S4.

The recovered time courses show a TOF-dependent response during the Sudoku task. The early gate at *τ*_*s*_ = 400 ps remained close to baseline, whereas the intermediate and late gates showed progressively larger increases in *αD*_*B*_(*t*). The strongest response was observed in the late gate at *τ*_*s*_ = 800 ps. Using the fitted baseline optical properties, with 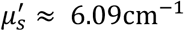, the *κ*^2^-derived activation index increased by +0.8 ± 1.9% at 400 ps, +9.8 ± 2.2% at 600 ps, and +15.2 ± 5.6% at 800 ps relative to baseline. This monotonic increase with TOF is consistent with stronger weighting of deeper tissue dynamics by later-arriving photons.

The phase-averaged *αD*_*B*_ (*τ*_*s*_) profiles further support this interpretation. During the Sudoku phase, *αD*_*B*_ increased primarily at later TOFs, whereas the early-TOF response remained small. The recovery phase remained partially elevated at late TOFs, suggesting a delayed return toward baseline after task completion. Because this is a single-subject proof-of-concept measurement, these observations should be interpreted as evidence of feasibility and depth-dependent sensitivity rather than as a quantitative estimate of prefrontal neurovascular activation.

The direct *g*_1_-fit analysis produced a qualitatively similar TOF-dependent activation trend, with activation indices of +1.56 ± 1.21% at 400 ps, +8.0 ± 2.15% at 600 ps, and +12.45 ± 2.99% at 800 ps, respectively, as shown in Fig. S5. Thus, both estimators support the same depth-dependent response pattern. However, the *κ*^2^-based recovery produced the larger late-gate activation amplitude and was used as the primary functional readout, consistent with the phantom and cuff-occlusion results showing improved stability of the temporal speckle-contrast representation in late TOF gates.

## 5. Discussion

We extended the iSCOS framework to the swept-source regime and demonstrated that temporal speckle-contrast analysis derived from the TOF-resolved field autocorrelation delivers depth-resolved, functional blood-flow information from a single-channel iNIRS platform. Three results support this conclusion. First, the simulated *κ*^2^(*τ*_*s*_, *T*) curves matched the analytical forward model [Eq. (12)] across *µ*_*s*_′, TOF gate, and exposure (Figs. 3–5), confirming that the TOF axis introduced by the wavelength sweep enters the speckle-contrast formalism without violating the underlying DWS assumptions. Second, the *g*_1_-based estimator was systematically more noise-robust than the direct estimator across the tested TOF gates, with the gap most pronounced at high noise amplitudes and in gates where the dynamic speckle contribution was weakest relative to the noise floor (Fig. 4). This advantage carried over to the simulated bi-layer *ξ*(*τ*_*s*_) recovery (Fig. 6). Third, the phantom and *in vivo* recordings (Figs. 7–10) show depth-selective dynamic information: homogeneous phantoms recover approximately constant flow index across TOF, bi-layer phantoms reproduce the predicted direction of the TOF-dependent *ξ*(*τ*_*s*_) slope transition, and the *in vivo* measurements show late-TOF amplification during vascular and cognitive activation paradigms.

Compared with our previous CW-iSCOS implementation [24], the present system retains the noise-robustness advantage of *g*_1_-derived speckle contrast and adds intrinsic depth selectivity at the cost of sequential wavelength tuning. Compared with TD-DCS [7,8] and SPAD-based parallel DCS [11], single-channel TOF-iSCOS shares the depth selectivity that motivates time-domain methods. Compared with TD-SCOS [31,32], our approach measures the optical field directly rather than its intensity autocorrelation, removing the Siegert-relation step and providing field-level noise correction. The trade-off is sweep-rate– limited temporal resolution per TOF volume. We discuss the underlying hardware bottleneck and our path forward in the next paragraph.

A key practical limitation of the present implementation is that detection runs through a single interferometric channel rather than a two-dimensional camera, which appears to forfeit the parallel-detection advantage that motivated CW-πNIRS and CW-iSCOS [13,24]. This is, however, not a fundamental property of TOF-iSCOS but a current hardware mismatch: TOF-resolved acquisition requires sampling the swept-source interferogram at digitizer rates on the order of 100 MHz per channel to preserve the optical-frequency bandwidth that defines the TOF axis, whereas the high-speed two-dimensional speckle cameras used in CW-iSCOS-class systems top out near 1 MHz full-frame readout, which is two orders of magnitude below what the wavelength sweep delivers. Parallel TOF-iSCOS therefore awaits either faster two-dimensional sensors or distributed/cropped-mode readout architectures (e.g., line-scan cameras synchronised to the sweep, region-of-interest streaming, on-chip pre-correlation, or multi-pixel SPAD arrays operated in a complex-field mode). Several of these approaches are under active development in our laboratory and elsewhere, and we view them as the natural follow-up to the single-channel demonstration reported here. In the meantime, single-channel TOF-iSCOS already offers two concrete advantages over single-channel TD-DCS [7,8]. First, the interferometric reference amplifies the weak diffuse sample field at the field level, restoring shot-noise-limited detection without requiring photon-counting hardware. Second, the field autocorrelation is measured directly, so no Siegert relation is invoked to convert from an intensity autocorrelation. This removes one source of systematic bias whenever the underlying speckle statistics deviate from circular-Gaussian [33] (e.g., for low photon counts, partially developed speckle, or strong static-scatterer contributions). The cost, already analysed in Section 2.4 and Section 4.2, is that single-channel detection cannot construct the direct spatial-variance estimator and must rely exclusively on the *g*_1_-derived *κ*^2^ route, which is precisely the estimator that the noise analysis identifies as preferable under additive sensor noise. As a result, the noise-robustness advantage demonstrated in CW-iSCOS [24] is preserved rather than lost in the migration to TOF-resolved acquisition.

The simulations in Fig. S1 further indicate how this limitation should scale with detector parallelism. As the number of independent detector pixels increases, the direct speckle-contrast estimator approaches the theoretical *κ*^2^ curve and the residual RMSE decreases, whereas the *g*_1_-derived estimator remains comparatively insensitive to detector pixel count because it obtains contrast from temporal autocorrelation rather than an instantaneous spatial ensemble. Thus, the present single-channel implementation is a deliberately conservative operating point: it demonstrates the TOF-resolved *κ*^2^ pipeline without the parallel speckle ensemble that should improve precision in future camera- or array-based systems.

Second, the in vivo demonstrations show that TOF-iSCOS can recover physiologically meaningful hemodynamic trends in both cuff-occlusion and cognitive-activation paradigms. During forearm occlusion, αDB decreased during cuff inflation and recovered after release, while preserving the expected TOF dependence. During the forehead Sudoku task, the *κ*^2^-derived response increased with TOF. The direct *g*_1_-fit analysis produced a similar late-gate amplification in the supplementary analysis, supporting the robustness of the observed depth trend.

Several limitations remain. The *in vivo αD*_*B*_ conversion relies on a homogeneous DWS model and on 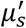 fitted from baseline TPSFs; residual optical-property errors therefore propagate directly into the absolute BFI scale. Layer-specific optical properties, partial-volume effects, and extracerebral contamination remain simplified in the present analysis. Future work should combine TOF-resolved contrast fitting with subject-specific optical-property estimation and multi-layer forward models, particularly for brain measurements where scalp and skull contributions are substantial.

Multiplicative noise (laser intensity, mechanical motion) affects both estimators. Future work should add a model term capturing it. Combined with the single-channel limitation discussed above, the absence of independent spatial speckle realizations means that the present demonstrations rely on temporal ensembles across consecutive sweeps. The parallel hardware roadmap outlined above will restore access to the direct spatial-variance estimator and enable single-shot speckle ensembles, which is a prerequisite for clinical-grade temporal resolution.

TOF-iSCOS sits at the intersection of interferometry (field-level access), time-domain methods (depth selectivity), and temporal speckle-contrast analysis (finite-exposure flow sensitivity). It converts a single swept-source iNIRS acquisition into a TOF-resolved, depth-selective, noise-corrected blood-flow estimate. The forearm and Sudoku recordings indicate feasibility for functional studies. Potential translation will require integrating sweep-rate–optimized swept-source hardware, real-time TOF reconstruction, motion-compensated probe designs, and cohort-level validation.

## 6. Conclusion

We have demonstrated time-of-flight–resolved interferometric speckle-contrast optical spectroscopy (TOF-iSCOS) as a single-channel proof-of-concept platform for depth-resolved functional blood-flow sensing. Swept-source iNIRS acquisition combined with *g*_1_-derived temporal speckle-contrast analysis at TOF gates yields *κ*^2^(*τ*_*s*_, *T*) curves that match DWS theory in simulations and homogeneous phantoms, reproduces the predicted bi-layer *ξ* trend in layered phantoms, and tracks depth-dependent hemodynamic responses in vivo during cuff occlusion and a Sudoku-task feasibility recording. Compared with non-gated CW-iSCOS, TOF-iSCOS adds depth selectivity while preserving the noise robustness of correlation-derived temporal speckle contrast. The present single-channel implementation is therefore a first methodological demonstration of TOF-resolved speckle-contrast analysis within swept-source iNIRS. Translation toward scalable functional monitoring will require multi-channel or camera-based detection, optimized sweep acquisition and processing, and cohort-level validation.

## Data availability

Data underlying the results presented in this paper are not publicly available at this time but may be obtained from the authors upon reasonable request.

## Supplementary Figures

The Supplementary Figures provide additional estimator comparisons, fitting diagnostics, and validation analyses supporting the main TOF-iSCOS results. They document the consistency between *g*_1_-based and *κ*^2^-based recovery, show the robustness of the temporal speckle-contrast estimator under noise and finite sampling, and provide supporting fits for the phantom and *in vivo* measurements presented in the main text.

**Figure S1.**
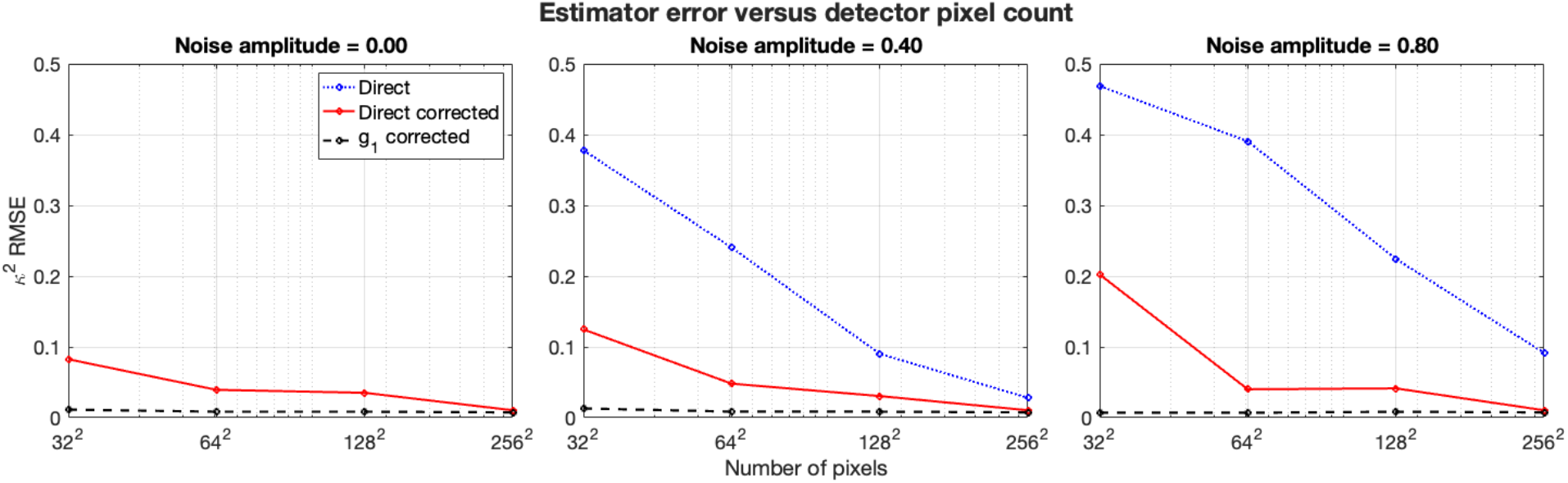
Influence of detector pixel count on estimator accuracy. The root-mean-square error (RMSE) between the estimated and theoretical *κ*^2^(*τ*_*s*_, *T*) curves was computed for detector arrays ranging from 32 × 32 to 256 × 256 pixels. Increasing detector parallelism substantially improves the accuracy of the direct estimator, indicating that finite speckle statistics are a dominant source of residual error. Standard variance subtraction further reduces the error but remains sensitive to detector size. In contrast, the corrected *g*_1_-based estimator exhibits consistently low error across all detector sizes, demonstrating that its noise robustness is largely independent of pixel count.

**Figure S2.**
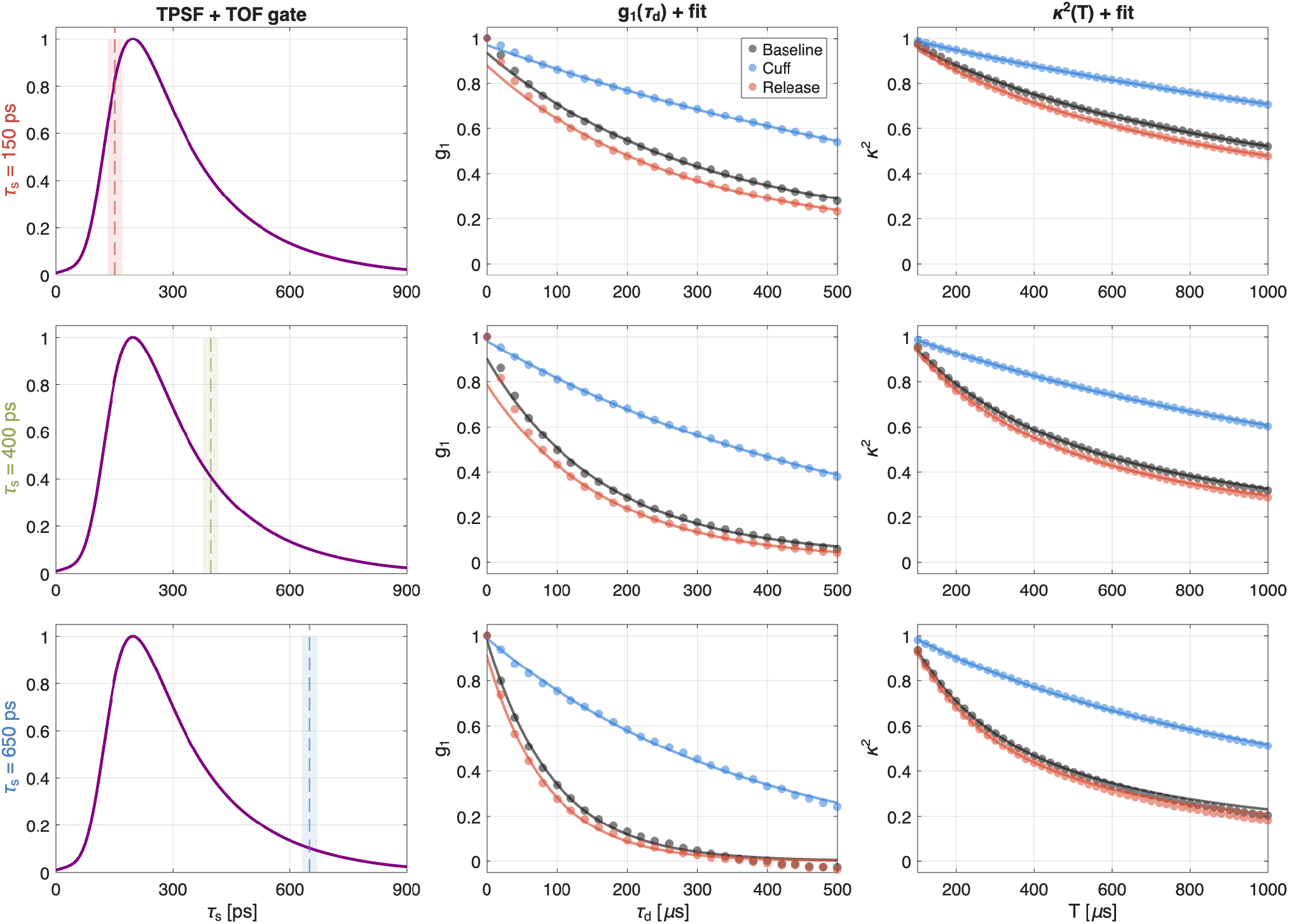
Forearm cuff-occlusion supporting fits. Rows show the representative TOF gates. Columns show the TPSF gate, the TOF-gated field autocorrelation *g*_1_(*τ*_*s*_, *τ*_*d*_) with fits, and the corresponding *g*_1_-derived temporal speckle-contrast curves *κ*^2^(*τ*_*s*_, *T*) with corresponding fits for the baseline, cuff, and release phases.

**Figure S3.**
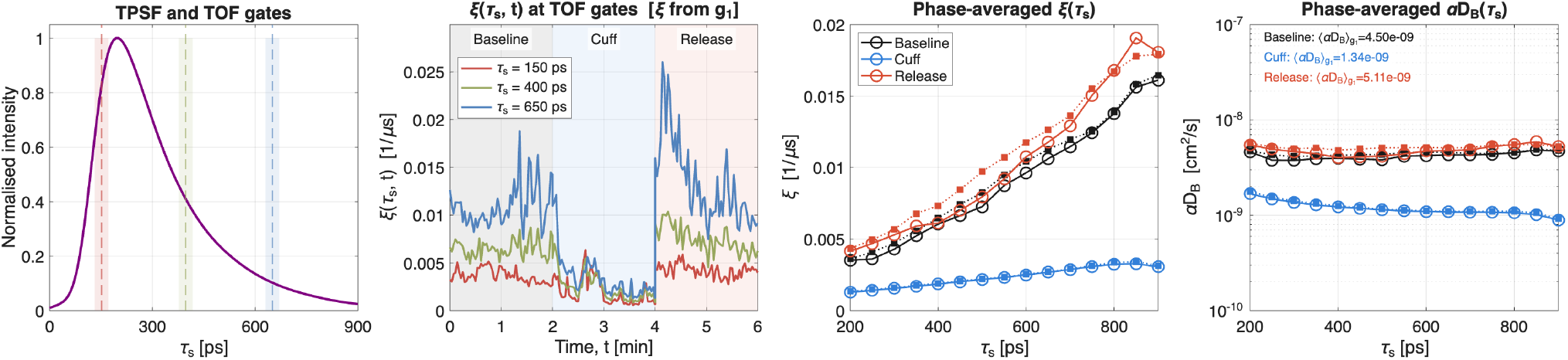
Forearm cuff-occlusion supporting analysis using direct *g*_1_-based recovery. The same TOF gates and physiological phases as in Fig. 9 are shown. The *g*_1_-based recovery reproduces the expected physiological response, with reduced decorrelation during cuff inflation and increased decorrelation during release. Compared with the primary *κ*^2^-based analysis, the recovered hyperemic peak amplitude is slightly lower, particularly in the late TOF gate, consistent with the lower stability of direct *g*_1_-based fitting in low-SNR late-photon regimes.

**Figure S4.**
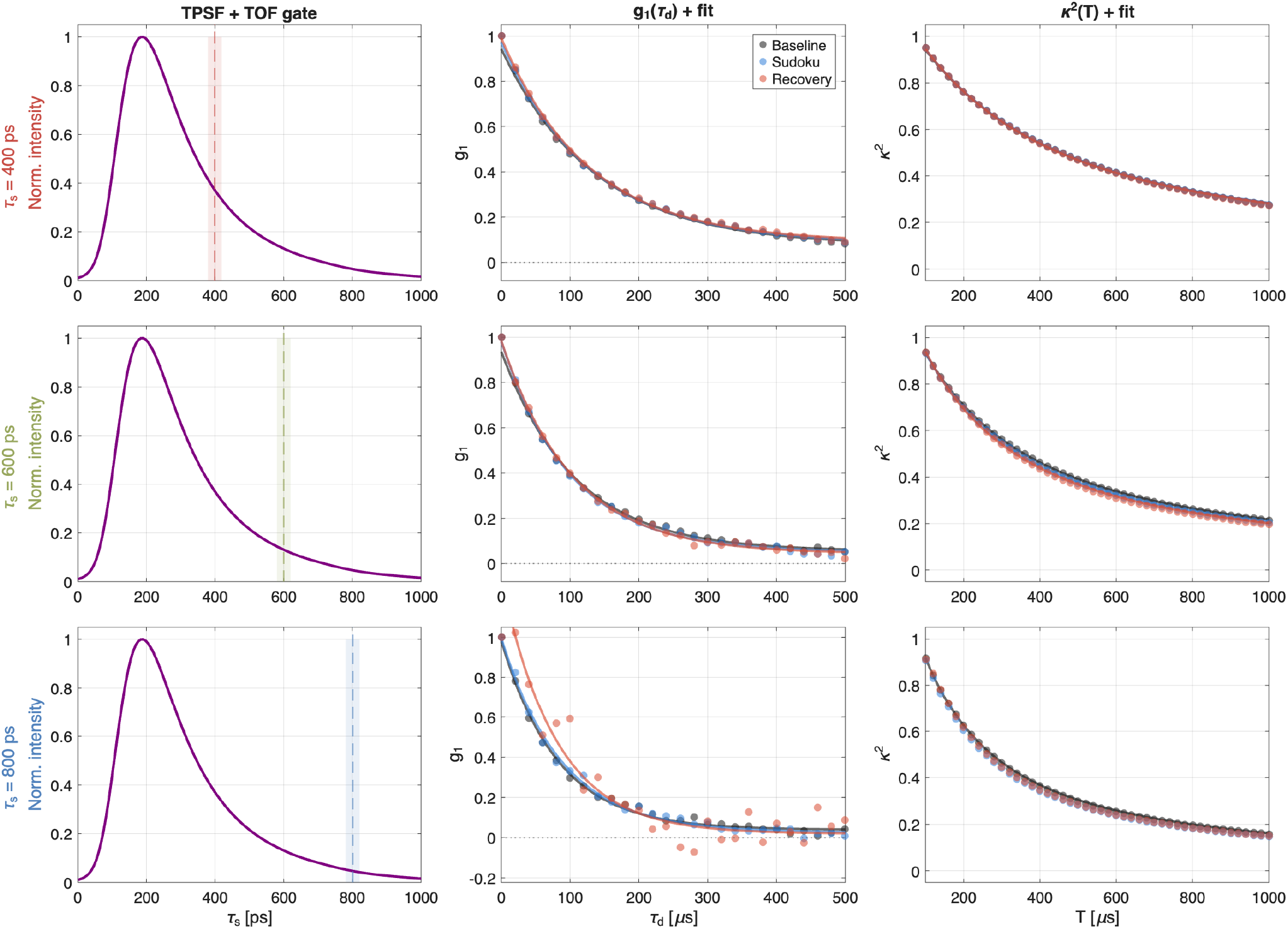
Forehead Sudoku supporting fits for the *κ*^2^-based analysis. Rows show the representative TOF gates *τ*_2_ = 400, 600, and 800 ps. Columns show the TPSF gate, the TOF-gated field autocorrelation *g*_1_(*τ*_*s*_, *τ*_*d*_ ) with fits, and the corresponding *g*_1_-derived temporal speckle-contrast curves *κ*^2^(*τ*_2_, *T*) with fits for the baseline, Sudoku, and recovery phases. The fits illustrate that the *κ*^2^-based recovery remains stable across the selected gates, including the late TOF gate used for the primary activation readout.

**Figure S5.**
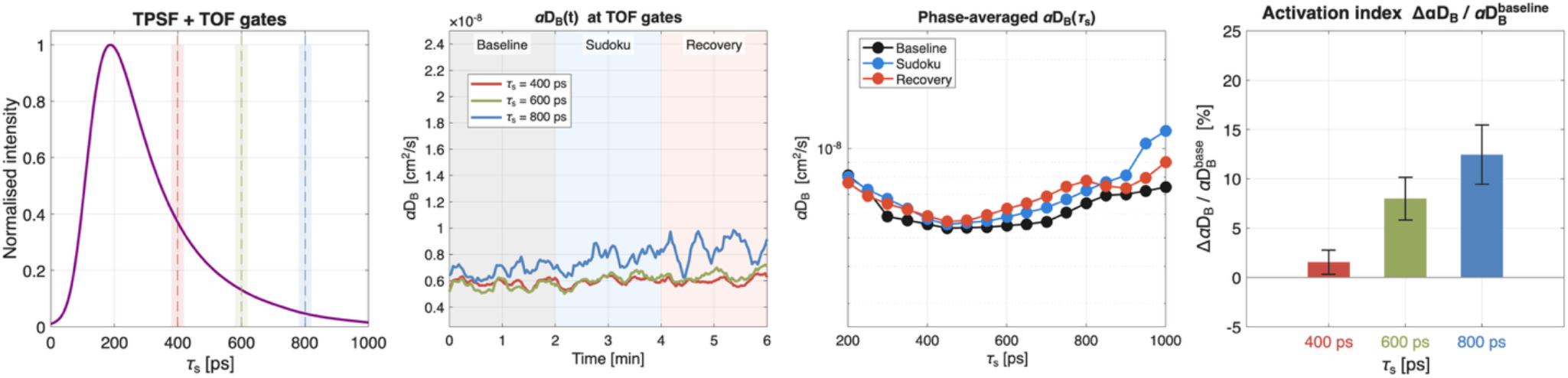
Direct *g*_1_-fit version of the forehead Sudoku analysis. The panels follow the same layout as the *κ*^2^-based analysis in Fig. 10. The direct *g*_1_-based recovery reproduces the same qualitative TOF-dependent activation trend, with minimal response in the early gate and progressively larger responses in the intermediate and late gates. The recovered late-gate activation amplitude is slightly lower than in the *κ*^2^-based analysis, supporting the use of *κ*^2^-based recovery as the primary functional TOF-iSCOS readout.

